# Core and indicative bacterial and fungal taxa define characteristic soil communities of arable land, grassland, and forest

**DOI:** 10.1101/2021.06.07.447343

**Authors:** Gschwend Florian, Hartmann Martin, Mayerhofer Johanna, Hug Anna, Enkerli Jürg, Gubler Andreas, Reto G. Meuli, Frey Beat, Widmer Franco

## Abstract

Soil microbial diversity has major influences on ecosystem functions and services. However, due to its complexity and uneven distribution of abundant and rare taxa, quantification of soil microbial diversity remains challenging and thereby impeding its integration into long-term monitoring programs. Using metabarcoding, we analyzed soil bacterial and fungal communities over five years at thirty long-term soil monitoring sites from the three land-use types, arable land, permanent grassland, and forest. Unlike soil microbial biomass and alpha-diversity, microbial community compositions and structures were site- and land-use-specific with CAP reclassification success rates of 100%. The temporally stable site core communities included 38.5% of bacterial and 33.1% of fungal OTUs covering 95.9% and 93.2% of relative abundances. We characterized bacterial and fungal core communities and their land-use associations at the family-level. In general, fungal families revealed stronger land-use type associations as compared to bacteria. This is likely due to a stronger vegetation effect on fungal core taxa, while bacterial core taxa were stronger related to soil properties. The assessment of core communities can be used to form cultivation-independent reference lists of microbial taxa, which may facilitate the development of microbial indicators for soil quality and the use of soil microbiota for long-term soil biomonitoring.

## 1. Introduction

Soil microorganisms constitute the majority of soil biodiversity (Bardgett and van der Putten 2014) and are main drivers of many soil processes (Costa *et al*. 2018, Hallin *et al*. 2018). A detailed understanding of belowground microbial diversity and of its influencing factors is the basis for a holistic view and understanding of ecosystem processes in terrestrial environments. However, a census of soil microorganisms remains largely incomplete, due to the enormous diversity and range of abundances of soil microorganisms. High microbial diversities have been observed at different scales ranging from aggregate (Hemkemeyer *et al*. 2019, Hemkemeyer *et al*. 2018), to landscape (Karimi *et al*. 2018), and global assessments (Bahram *et al*. 2018, Větrovský *et al*. 2019).

At the land-scape scale, soil bacterial and fungal diversities are strongly correlated to soil pH (Griffiths *et al*. 2011, Lauber *et al*. 2009), which is caused by direct effects but also by indirect effects such as changing the availability of nutrients (Glassman *et al*. 2017, Lammel *et al*. 2018). The number of bacterial taxa in soils depends on the pH and has been reported to reach its maximum at pH values between 6 and 7 (Lauber *et al*. 2009). Furthermore, community structures of soil bacteria change with pH, because specific bacterial taxa reveal distinct pH preferences. For instance, within the phylum Acidobacteria, taxa belonging to the class Acidobacteriia are in general negatively correlated to soil pH, while taxa belonging to Acidobacteria Subgroup 6 commonly reveal a positive correlation to soil pH (Kielak *et al*. 2016). Further drivers of bacterial community structures depend on the system studied and include factors such as soil texture, climate, and plant communities (Bahram *et al*. 2018, Griffiths *et al*. 2016, Karimi *et al*. 2018, Leff *et al*. 2018). In comparison to soil bacterial diversity, soil fungal diversity has been shown to be geographically more structured (Bahram *et al*. 2018, Talbot *et al*. 2014). In a global meta-analysis that covered 742 sites, Větrovský *et al*. (2019) identified climate factors as main drivers of soil fungal communities, followed by soil properties, and vegetation parameters. Finally, factors related to land management, such as agricultural intensity (Banerjee *et al*. 2019), tillage (Babin *et al*. 2019, Degrune *et al*. 2017), fertilization (Hartmann *et al*. 2015, Piazza *et al*. 2019), or compaction (Hartmann *et al*. 2014) may influence diversity of soil bacteria and fungi. While the major environmental determinants of soil bacterial and fungal communities are largely known, less is known about common components of these communities, their taxonomic representatives, and their diversities.

Surveys of soil bacterial and fungal communities usually reveal a large number of unknown taxa. Delgado-Baquerizo (2019) has reported that in a global survey 99% of bacterial and 63% of fungal OTUs remained unclassified at the species-level, and that the number of unclassified bacterial or fungal OTUs at the phylum-level in a sample has ranged between 1.4% and 9.4%. In a meta-analysis on the global diversity of soil fungi, an average of only 53% of the sequences per sample could be assigned to entries in the UNITE reference database, which notably includes sequences from environmental samples (Větrovský *et al*. 2019). High ratios of unclassified sequences at the species level may be due to a lack of resolution of the used DNA barcodes (e.g. Gschwend *et al*. 2021), or due to missing reference sequences. To elucidate the unknown microbial diversity and describe consistently occurring OTUs, several attempts have been made to identify the most common taxa, which could constitute a core of soil microbial communities (Delgado-Baquerizo *et al*. 2018, Egidi *et al*. 2019). OTUs contributing to the global bacterial soil core community were assigned in descending order of relative abundance to the phyla Proteobacteria, Actinobacteria, Planctomycetes, Chloroflexi, Verrucomicrobia, Bacteroidetes, Gemmatimonadetes, Firmicutes, Armatimonadetes, Saccharibacteria, and candidate division WS2 (Delgado-Baquerizo *et al*. 2018). Five of these phyla, i.e., Proteobacteria, Actinobacteria, Planctomycetes, Bacteroidetes, and Firmicutes, have also been reported among those with an average relative abundance of at least 5% in a soil bacterial survey across France (Karimi *et al*. 2018), which has identified Acidobacteria as an additional dominant phylum. Dominant soil bacterial phyla have revealed distinct ecological preferences such as Alphaproteobacteria and Verrucomicrobia that were more abundant in forest and permanent grassland as compared to arable and vineyard soils, while the inverse was found for Chloroflexi and Gemmatimonadetes (Karimi *et al*. 2018). However, diverse habitat associations are often detected for taxa assigned to the same phylum. For instance within the phylum Chloroflexi, the family Anaerolineaceae were associated to soils with pH above 5, while Ktedonobacteraceae were associated to a lower soil pH (Mayerhofer *et al*. 2021). For soil fungi, a global survey of 365 sites has revealed Ascomycota, Basidiomycota, Mortierellomycota, and Mucoromycota as dominant fungal phyla in soils (Tedersoo et al. 2014), which has been largely confirmed, although the high abundance of Mortierellomycota has been questioned (Větrovský *et al*. 2019). Egidi *et al*. (2019) have proposed that globally dominant soil fungal OTUs almost exclusively derived from Ascomycota with 80 of 83 dominant fungal OTUs classified to this phylum. Despite the recent interest in taxonomic surveys of soil bacterial (Delgado-Baquerizo *et al*. 2018, Karimi *et al*. 2018, Walsh *et al*. 2019) and fungal diversity (Egidi *et al*. 2019, Tedersoo *et al*. 2017), habitat associations of soil bacteria and fungi at lower taxonomic levels are still largely lacking.

In a previous study, thirty long-term monitoring sites of the Swiss Soil Monitoring Network (NABO) were surveyed over five years, and it has been shown that soil bacterial and fungal communities of different sites remained temporally stable and compositionally distinct (Gschwend *et al*. submitted). However, that study has focused on community structures and treated OTUs as anonymous entities without assessing their taxonomy. Furthermore, temporal dynamics of soil bacterial and fungal community structures have been assessed but detailed analyses of environmental drivers of community structures among land-use types, and the individual sites have not been provided. Detailed descriptions of habitat associations of bacterial and fungal taxa are, for instance, also needed to develop microbial indicators for biological assessments of soil quality.

Here, we assess bacterial and fungal diversity and community structures at thirty sites of the NABO. Our main research goals were to characterize consistently detected OTUs over several years, which allow for a robust assessment of soil microbial communities along with their habitat associations. Specifically, our research aimed to i) assess site- and land-use-specific soil microbial communities; ii) identify OTUs, which are consistently detected (core OTUs) as well as taxa indicative of environmental factors (indicative OTUs); iii) assess the main environmental factors structuring core communities; iv) describe diversity and identity of core OTUs as well as their distribution among land-use types.

## 2. Material and Methods

### 2.1 Sampling design, DNA extraction, and microbial biomass measurement

Samples were taken during five years, from 2012 to 2016, at thirty sites (Figure S1) of the Swiss Soil Monitoring Network (NABO) in early spring after snow melt and before fertilization. Three land-use types, i.e. arable land, permanent grassland, and forests were sampled with ten sites each. Arable sites were managed with crop rotations, which included three to six different crops, and with one exception they were conventionally tilled. Forest sites included four coniferous, two mixed, and four deciduous forests. At each site, three composite samples composed of 25 soil cores of 20 cm depth and 2.5 cm diameter were taken from a 10 m by 10 m plot according to the standardized sampling protocol of the Swiss Soil Monitoring Network (Gubler *et al*. 2019). Samples were immediately stored at 4°C after sampling and processed within 48 hours. Homogenized soil was mixed with DNA extraction buffer ([2% hexadecyl trimethyl ammonium chloride (CTAB); 20 mM EDTA pH 8; 2 M NaCl; 100 mM tris hydroxymethylaminomethane pH 8; 2% polyvinylpyrrolidone (PVP-40)], (Lazzaro *et al*. 2006). Quantitative DNA extraction was achieved by extracting DNA three times from each sample following Bürgmann *et al*. (2001) with the modifications by Hartmann *et al*. (2005). DNA quantity was determined using PicoGreen (Invitrogen, Carlsbad, CA) on a Cary Eclipse fluorescence spectrophotometer (Varian, Inc. Palo Alto, CA) and cross-validated using Qubit 1.0 (Life Technologies, Carlsbad, CA, USA). DNA was cleaned using the NucleoSpin® gDNA clean-up kit (Machery-Nagel, Düren, Germany) according to the manufacturer’s instruction. Microbial biomass carbon (C_mic_) was assessed using chloroform-fumigation-extraction according to Vance *et al*. (1987) with a k_EC_ value of 0.45 (Joergensen 1996). Measurements of soil physico-chemical properties, i.e., soil pH, total and organic carbon, total nitrogen, C/N-ratio, bulk density, soil texture, and gravimetric water content, have been described in Gschwend et al. (submitted).

### 2.2 Barcode amplification, sequencing, and sequence analysis

Bacterial variable region 3 and 4 of the small sub-unit of the ribosomal RNA gene (16S rRNA) were amplified using primers 341F (5’ CCTAYGGGDBGCWSCAG 3’) and 806R (5’ GGACTACNVGGGTHTCTAAT 3’) (Frey *et al*. 2016). Fungal internal transcribed spacer 2 (ITS2) was amplified using primers ITS3 (5’ CAHCGATGAAGAACGYRG 3’) and ITS4 (5’ TCCTSCGCTTATTGATATGC 3’) (Tedersoo *et al*. 2014). Four reactions using the GoTaq® Hot Start Polymerase (Promega) were done for each sample using 20 ng of DNA for each reaction. Reactions were performed according to Mayerhofer *et al*. (2017) with two modifications, which were an initial denaturation at 95°C for two minutes, as well as 35 PCR cycles for the bacterial and fungal markers. Production of sequencing libraries and paired-end sequencing on an Illumina MiSeq v3 were performed at the Génome Québéc Innovation Center at the McGill University (Montréal, Canada). Raw sequences, (NCBI SRA: XXXXXX) were quality filtered using a custom sequence analysis pipeline largely based on USEARCH version 9 (Edgar 2010, Frey *et al*. 2016) and is described in greater detail in Gschwend et al. submitted). Only sequences occurring in at least two samples were allowed to form OTU centroids. Sequences were clustered into OTUs based on a 97% sequence identity threshold. This threshold was chosen to obtain a conservative estimate of soil microbial diversity and because diversity patterns between OTUs and sequence variants based approaches are highly correlated (Glassman and Martiny 2018). Taxonomic assignment was obtained using the RDP classifier implemented in mothur version 1.36.1 (Schloss *et al*. 2009) and a minimum bootstrap value of 80% with the SILVA 132 database (Quast *et al*. 2012) as reference for bacterial sequences. Eukaryotic sequences were classified with the same approach to a Genbank database (Frey *et al*. 2016) to discriminate between fungal and other eukaryote sequences. Fungal sequences were subsequently compared to the UNITE v 7.2 reference database (Nilsson *et al*. 2018).

### 2.3 Statistics

All analyses unless stated otherwise, were performed in R (R Core Team 2016, RStudio 2015). Mean values of environmental factors were calculated for samples taken at the same time point to avoid pseudo-replication. Similarly, calculations of alpha- and beta-diversity values were based on median values of OTUs per sampling time point. To get independent samples for the assessment of similarities and differences between land-use types, median values of OTUs were obtained per site followed by Jaccard and Bray-Curtis similarity calculations. Spearman correlations were used to link univariate responses to environmental factors. Multivariate responses of communities were assessed by PERMDISP (Anderson *et al*. 2006) to evaluate homogeneity of dispersions between groups and permutational analysis of variance (PERMANOVA, Anderson 2001) to analyse between group differences. PRIMER7 (Anderson *et al*. 2008, Clarke and Warwick 2001) was used for PERMANOVA. PERMANOVA design included land-use types as a fixed factor, sites as random factor nested within land-use type, and year as a random factor. Effects on community structures were expressed as square root of component of variation (√CV), which are in the unit of the original community dissimilarity, i.e., Bray-Curtis dissimilarity. The order of covariates in sequential PERMANOVA tests were selected based on the model selection algorithm implemented in distance-based linear model (DISTLM, McArdle and Anderson 2001) within PRIMER7, where AICc was chosen as model selection criterion. P-values of multiple tests were adjusted using Benjamini-Hochberg procedure (Benjamini and Hochberg 1995). Site specificity was further assessed by leave-one-out cross-validation based on linear discriminant analysis (LDA) for univariate and based on canonical analysis of principal coordinates (CAP, Anderson and Willis 2003) for community structures. LDA and CAP were calculated within R using the functions ‘train’ of the package caret (Kuhn 2008) and ‘CAPdiscrim’ of the package ‘BiodiversityR’ (Kindt and Coe 2005), respectively. Ternary plots were drawn using the R package ggtern 3.0.0.1 (Hamilton and Ferry 2018).

### 2.4 Definition of OTU groups

We distinguished two OTU groups, i.e., ‘core’ and ‘‘indicative’ OTUs, which included two or three subgroups, respectively (Table 1). Core OTUs were defined based on their consistent occurrence at a site or in a land-use type. Site core OTUs (sc-OTUs), were defined as OTUs that occur in at least 80% of the 15 samples from a given site. Similarly, land-use type core OTUs (lc-OTUs), were defined as OTUs that are sc-OTUs in at least 80% of the 10 sites of a given land-use type. Indicative OTUs included three subgroups, which were i) correlated to an environmental factor, ii) indicative for land-use types, or iii) indicative for an individual site. The first subgroup was defined based on a Spearman correlation of |rho| > 0.4 (p < 0.05) with an environmental factor. Subgroups two and three were defined based on indicator species analysis using the ‘indicspecies’ R-package (De Cáceres and Legendre 2009). OTUs with an adjusted p-value smaller than 0.05 and an indicator value higher than 0.8 for a single or a combination of land-use types, or for individual sites were termed ‘land-use-indicative’ and ‘site-indicative’ OTUs (Table 1). Therefore, land-use- and site-indicative OTUs have a significantly higher relative abundance and occurrence in a given land-use type or site. In contrast, the definition of core OTUs does not include information of the OTU abundance and occurrence in other land-use types or sites.

**Table 1.**
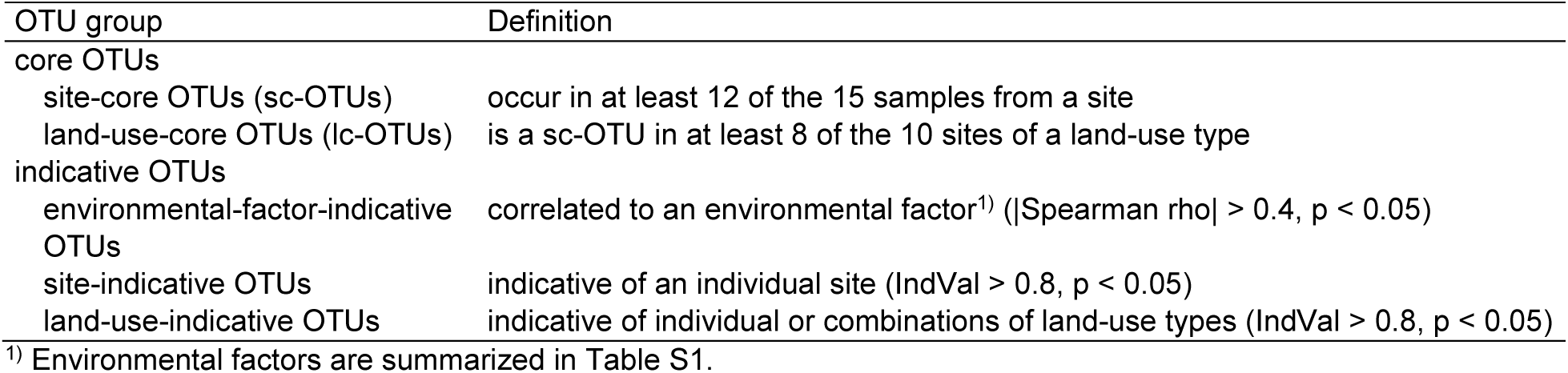
Definitions of OTU groups and subgroups (see also Table 3)

## 3. Results

### 3.1 Increasing resolution from microbial biomass to community structures

Thirty sites from three land-use types, i.e., ten each from arable land, permanent grassland, and forest, were surveyed with yearly samplings during five years, which yielded 450 samples. Soil microbial communities were assessed using three different approaches, which were i) soil microbial biomass, i.e., based on soil microbial carbon (C_mic_) content determined with chloroform fumigation extraction, and soil DNA content, that correlated (rho = 0.79, p < 0.0001), ii) alpha-diversity based on OTU richness, Simpson evenness, and inverse Simpson index, and iii) beta-diversity based on Jaccard similarities and Bray-Curtis dissimilarities (Table 2, but see also supplementary results for a summary of the sequencing data). Microbial biomass and alpha-diversity revealed no site- (reclassification ≤ 4.7%), and low land-use-specificity (reclassification ≤ 61.3%, Table 2). Values of both microbial biomass measures were significantly reduced in arable land (Tukey HSD, p ≤ 0.0007, Table S1), while bacterial alpha-diversity was increased in arable land (Tukey HSD, p = 0.0096, Table S1). Fungal alpha-diversity with the exception of fungal OTU richness were significantly lower in forest soils (Tukey HSD, p ≤ 0.01, Table S1). Community compositions (Jaccard similarity) and structures (Bray-Curtis dissimilarity) were land-use- (Figure 1) and site-specific with reclassification success rates of 100% for bacteria and fungi (Table 2). Consequently, information on community composition or structure was needed for resolving the different drivers of bacterial and fungal communities in soil.

**Table 2.**
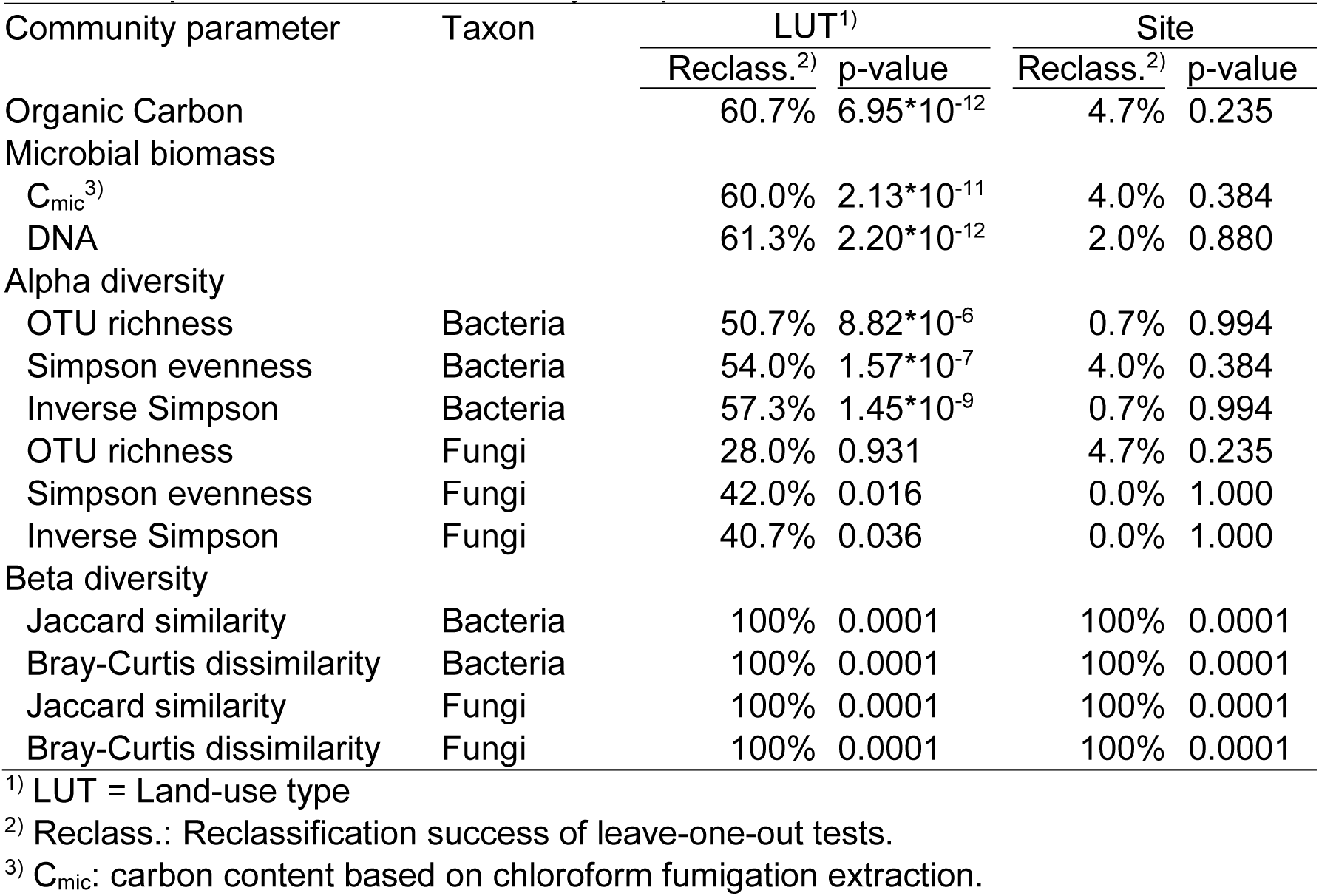
Site and land-use specific soil microbial communities at different analytical levels. Site and land-use type specificity was calculated using a leave-one-out reclassification test based on linear discriminant analysis for univariate and canonical analysis of principal coordinates (CAP) with 9 999 permutations for community compositions and structures.

**Figure 1:**
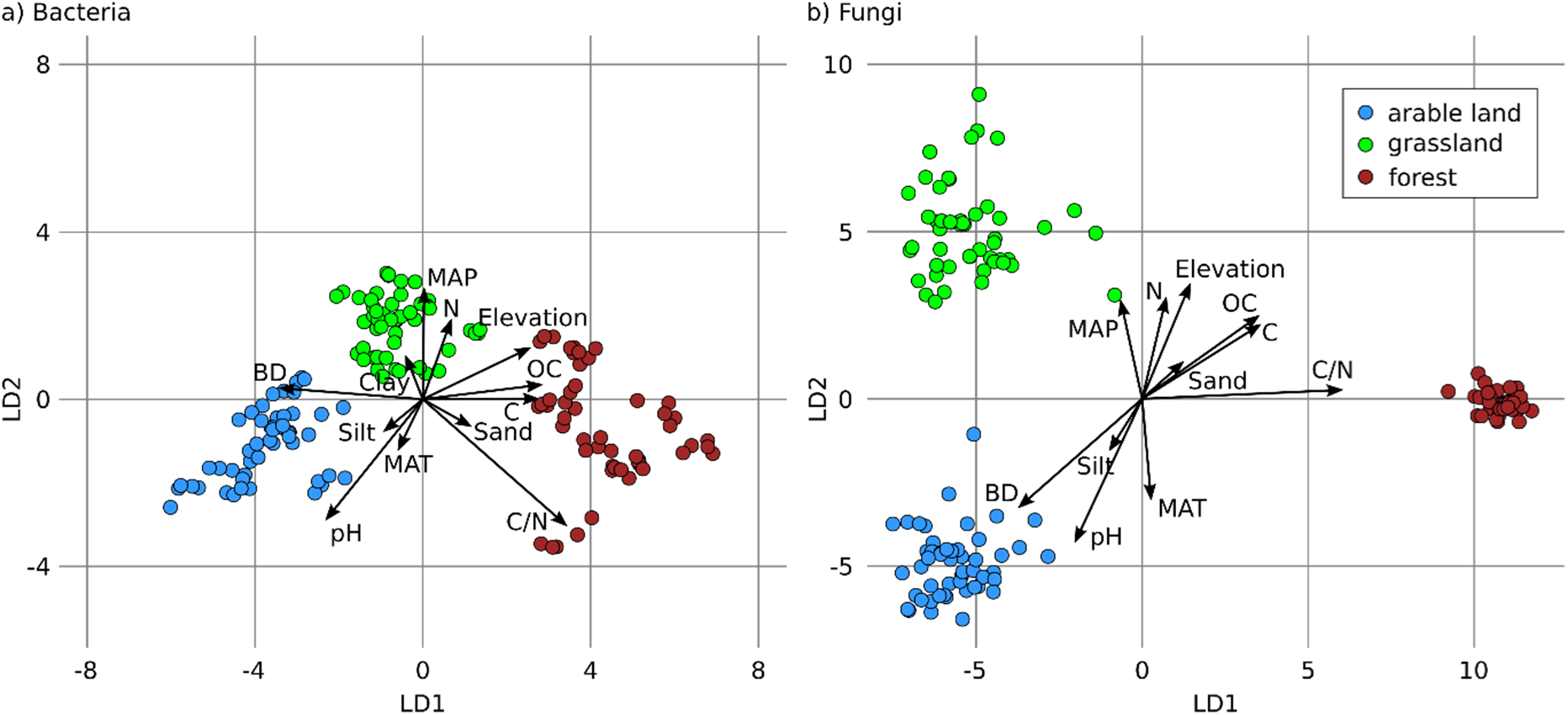
Separation of bacterial (A) and fungal (B) communities by land-use and correlated environmental factors. Three land-use types, i.e., arable land (blue), permanent grassland (green), and forest (brown), were sampled with 10 sites each. Per site, 15 samples were obtained with yearly triplicates during five years. Average communities for yearly replicates are shown (N = 150). Ordinations are based on canonical analyses of principal coordinates (CAP) constrained by land-use types. Axes show linear discriminants (LD). Arrows indicate significant correlations of communities to environmental factors, i.e., bulk density (BD), clay, silt, sand, pH, mean annual temperature (MAT), mean annual precipitation (MAP), ratio of C/N (C/N), total carbon (C) and nitrogen (N), organic carbon (OC), and elevation.

### 3.2 Partitioning of OTUs into core and indicative groups

The high site-specificity of soil bacterial and fungal community structures, which was maintained over five years, also reflected a high temporal stability. Temporally stable core taxa, i.e., site-core (sc) OTUs and land-use type core (lc) OTUs were defined as outlined in Table 1. Of the 18 140 bacterial OTUs (bOTUs) 6 979 (38.5%), which covered 95.9% relative abundance were classified as sc-OTUs and 1 136 of these sc-OTUs (covering 69.1% relative abundance) were also classified as lc-OTUs (Table 3). A similar proportion of the 8 477 fungal OTUs (fOTUs), i.e., 2 802 fOTUs (33.1%) and covering 93.2% relative abundance, was classified as sc-OTUs, but only 103 of them (29.4% relative abundance) were also classified as lc-OTUs. In addition to these core taxa, we defined indicative OTUs, i.e., OTUs that structured communities according to environmental conditions. More specifically, we distinguished three categories of indicative OTUs, i.e., OTUs correlated to environmental factor, as well as OTUs indicative for land-use types and OTUs indicative of a given site (see Table 1 for definitions). Most strikingly, the number and particularly the abundance of site-indicative OTUs was higher for fungi (1 445 fOTUs, 29.9% relative abundance), as compared to bacteria (1 146 bOTUs, 3.1% relative abundance). The vast majority of indicative OTUs were also classified as sc-OTUs (95% for bacteria, 90% for fungi, Figure S2). Communities composed of only sc-OTUs, i.e., core communities, were almost perfectly correlated (rho ≥ 0.97) to the entire communities, both in terms of alpha- and beta-diversity (Table S2). Consequently, soil microbial core communities are representative of the respective entire communities. The following analyses were therefore based on these core communities.

**Table 3.** Summary of OTU partitioning into core and indicative groups and subgroups. Core and indicative OTUs were defined at the site and the land-use type level (see Table 1 for definitions of OTU groups and subgroups). LUT = land-use type.

|  | OTU group | Subgroup | OTUs [N] | Abundance [%] | Correlation<sup>1)</sup> [rho] | Phyla [N] | Families [N] |
| --- | --- | --- | --- | --- | --- | --- | --- |
| Bacteria | Core | Site | 6 979 | 95.9 | 1.000 | 31 | 215 |
|  | Core | LUT | 1 136 | 69.1 | 0.995 | 17 | 119 |
|  | indicative | Environmental factor | 3 103 | 67.0 | 0.983 | 27 | 164 |
|  | indicative | Site | 1 146 | 3.1 | 0.736 | 28 | 106 |
|  | indicative | LUT | 699 | 27.2 | 0.931 | 17 | 102 |
|  | All |  | 18 140 | 100 |  | 46 | 320 |
| Fungi | Core | Site | 2 802 | 93.2 | 0.999 | 9 | 176 |
|  | Core | LUT | 103 | 29.4 | 0.893 | 5 | 35 |
|  | indicative | Environmental factor | 553 | 42.5 | 0.942 | 7 | 96 |
|  | indicative | Site | 1 445 | 29.9 | 0.765 | 7 | 125 |
|  | indicative | LUT | 171 | 35.2 | 0.891 | 5 | 50 |
|  | All |  | 8 477 | 100 |  | 12 | 304 |

<sup>1)</sup> Spearman correlation to entire community (Mantel test)

### 3.3 Environmental factors driving structures of core communities

Soil bacterial and fungal core communities were mainly structured by soil pH and the C/N-ratio (Table 4). In addition to the environmental factors considered, land-use type and site significantly explained variance of soil bacterial (√CV_Land-use type_ = 0.23, √CV_Site_ = 0.31), and fungal (√CV_Land-use type_ = 0.31, √CV_Site_ = 0.49) community structures. Soil pH was the strongest driver for bacterial community structures overall and within each land-use type (Table S3). The second strongest environmental factor in the overall analysis was the C/N-ratio, but it had no or minimal effects on the community structures within land-use types (Table S3 & S4). This may be due to the clear difference in C/N-ratio between forest and the other two land-use types (Table S1), indicating that a high C/N-ratio represented a proxy for forest soils in the overall analysis. The separate analysis of arable sites also allowed to consider crop as an additional factor shaping microbial communities (Table S3 & S4), which was more strongly affecting fungal (√CV = 0.16) as compared to bacterial (√CV = 0.06) core communities. In line with the data on core community structures, the strongest correlations of individual OTUs to environmental factors were detected with soil pH for bacterial OTUs (Table S5), and with soil pH, C/N-ratio, and organic carbon for fungal OTUs (Table S6).

**Table 4.** Effects of environmental factors on bacterial (A) and fungal (B) communities as assessed by PERMANOVA. Factors are sorted by their position in the PERMANOVA model with environmental factors as covariates. Year and site were random factors with site being nested within land-use type. Factors below the dotted lines are categorical. Significance codes: *** p < 0.001, ** p < 0.01, * p < 0.05

a) Bacteria
| Env. factor<sup>1)</sup> | Pseudo-F | $\sqrt{CV}^{2)}$ | p-value | |
| --- | --- | --- | --- | --- |
| pH | 18.6 | 0.25 | 0.0001 | *** |
| C/N-ratio | 5.4 | 0.15 | 0.0001 | *** |
| MAP<sup>3)</sup> | 2.2 | 0.07 | 0.0001 | *** |
| Clay | 2.0 | 0.06 | 0.0001 | *** |
| Elevation | 1.4 | 0.05 | 0.0119 | * |
| C<sub>org</sub> | 1.5 | 0.07 | 0.0130 | * |
| Sand | 1.3 | 0.06 | 0.0338 | * |
| MAT<sup>4)</sup> | 1.2 | 0.04 | 0.1418 |  |
| Year | 4.2 | 0.05 | 0.0001 | *** |
| LUT<sup>5)</sup> | 2.9 | 0.23 | 0.0017 | ** |
| Site | 19.4 | 0.31 | 0.0001 | *** |
| LUT<sup>5)</sup> x Year | 1.5 | 0.04 | 0.0001 | *** |
| Residuals |  | 0.15 |  |  |

b) Fungi
| Env. factor<sup>1)</sup> | Pseudo-F | $\sqrt{CV}^{2)}$ | p-value | |
| --- | --- | --- | --- | --- |
| C/N-ratio | 5.5 | 0.19 | 0.0001 | *** |
| pH | 2.6 | 0.14 | 0.0001 | *** |
| Elevation | 1.5 | 0.09 | 0.0086 | ** |
| Sand | 1.3 | 0.06 | 0.0330 | * |
| Clay | 1.2 | 0.06 | 0.1399 |  |
| MAT<sup>4)</sup> | 1.2 | 0.06 | 0.1012 |  |
| C<sub>org</sub> | 1.2 | 0.06 | 0.1226 |  |
| Bulk density | 1.3 | 0.08 | 0.1061 |  |
| Year | 2.2 | 0.06 | 0.0001 | *** |
| LUT<sup>5)</sup> | 2.5 | 0.31 | 0.0001 | *** |
| Site | 14.4 | 0.49 | 0.0001 | *** |
| LUT<sup>5)</sup> x Year | 1.3 | 0.05 | 0.0001 | *** |
| Residuals |  | 0.28 |  |  |

<sup>1)</sup> Env. factor: environmental factor;
<sup>2)</sup> $\sqrt{CV}$ : square root of component of variation, expressed as Bray-Curtis dissimilarity;
<sup>3)</sup> MAP: mean annual precipitation
<sup>4)</sup> MAT: mean annual temperature
<sup>5)</sup> LUT: Land-use type

### 3.4 Association of bacterial and fungal core OTUs to land-use types

The similarities of bacterial and fungal communities among land-use types were highest between arable and permanent grassland soils, while they were lowest between arable and forest soils (AG and AF in Figure 2). The similarity between communities from forest and permanent grassland sites was higher for bacteria than for fungi, which was particularly striking, when relative abundances were considered as accounted for in Bray Curtis similarities (FG in Figure 2c & d). To assess these differences in greater detail, the distribution of core taxa among the land-use types were analyzed using ternary plots, which depict the abundance of sc-OTUs in each land-use type and in all combinations (Figure 3). The ternary plots clearly revealed different distributions of bacterial and fungal sc-OTUs among land-use types. On the one hand, bacterial sc-OTUs were distributed among the land-use types and all their combinations except for the combination of ‘arable land and forest’, for which only two lc-OTUs were detected (Figure 3a). Eighty-seven bacterial sc-OTUs were core of all three land-use types (AGF in Figure 3a). On the other hand, fungal sc-OTUs were accumulated along the axes of arable land to permanent grassland and in forest (Figure 3b). Only three fungal sc-OTUs were cores of all three land-use types and no land-use type core was detected for the combination of arable land and forest (Figure 3b). The difference in bacterial and fungal distributions among the land-use types was also evident from the number of sc-OTUs with at least 80% of their abundance in a single land-use type (Figure 3, red tips of the ternary plots). For bacteria, the number of such sc-OTUs that not necessarily represented an lc-OTU, was highest in arable soils (1 239 bOTUs), slightly less in forest (967 bOTUs), and lowest in permanent grassland (308 bOTUs). For fungi, more sc-OTUs were predominantly detected in forest (1 231 fOTUs), as compared to permanent grassland (502 fOTUs) and arable land (424 fOTUs).

**Figure 2:**
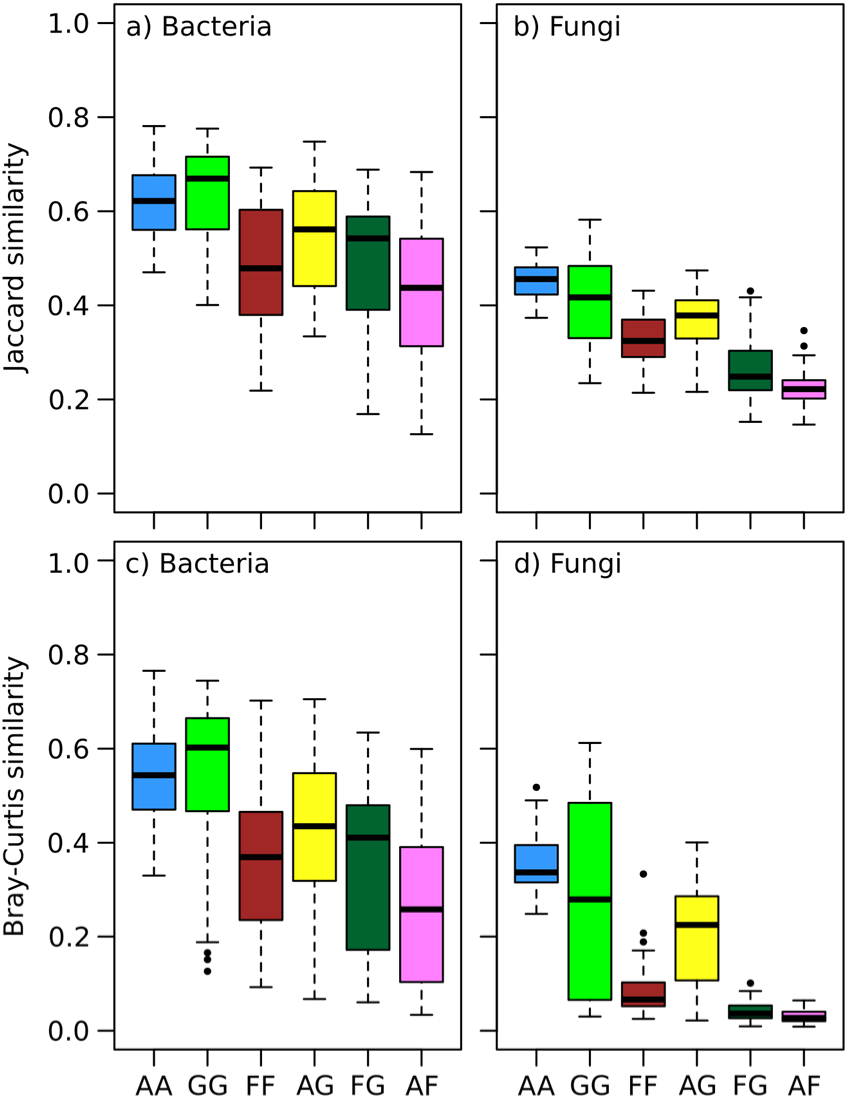
Pairwise comparisons of bacterial (a, c) and fungal (b, d) communities composed of core OTUs for a site, i.e., OTUs that occurred in at least 12 of the 15 samples from a site. Boxplots showing Jaccard (a, b) and Bray-Curtis (c, d) similarities between two sites depending on their land-use type. The Jaccard similarity corresponds to the ratio of shared OTUs between two sites, while the Bray-Curtis similarity takes also the relative abundance of each OTU into account. Sites of three land-use types, i.e. arable land (A), grassland (G), and forest (F), were assessed in pairwise combinations of the same land-use type (AA, GG and FF) as well as between different land-use types (AG, FG and AF).

**Figure 3:**
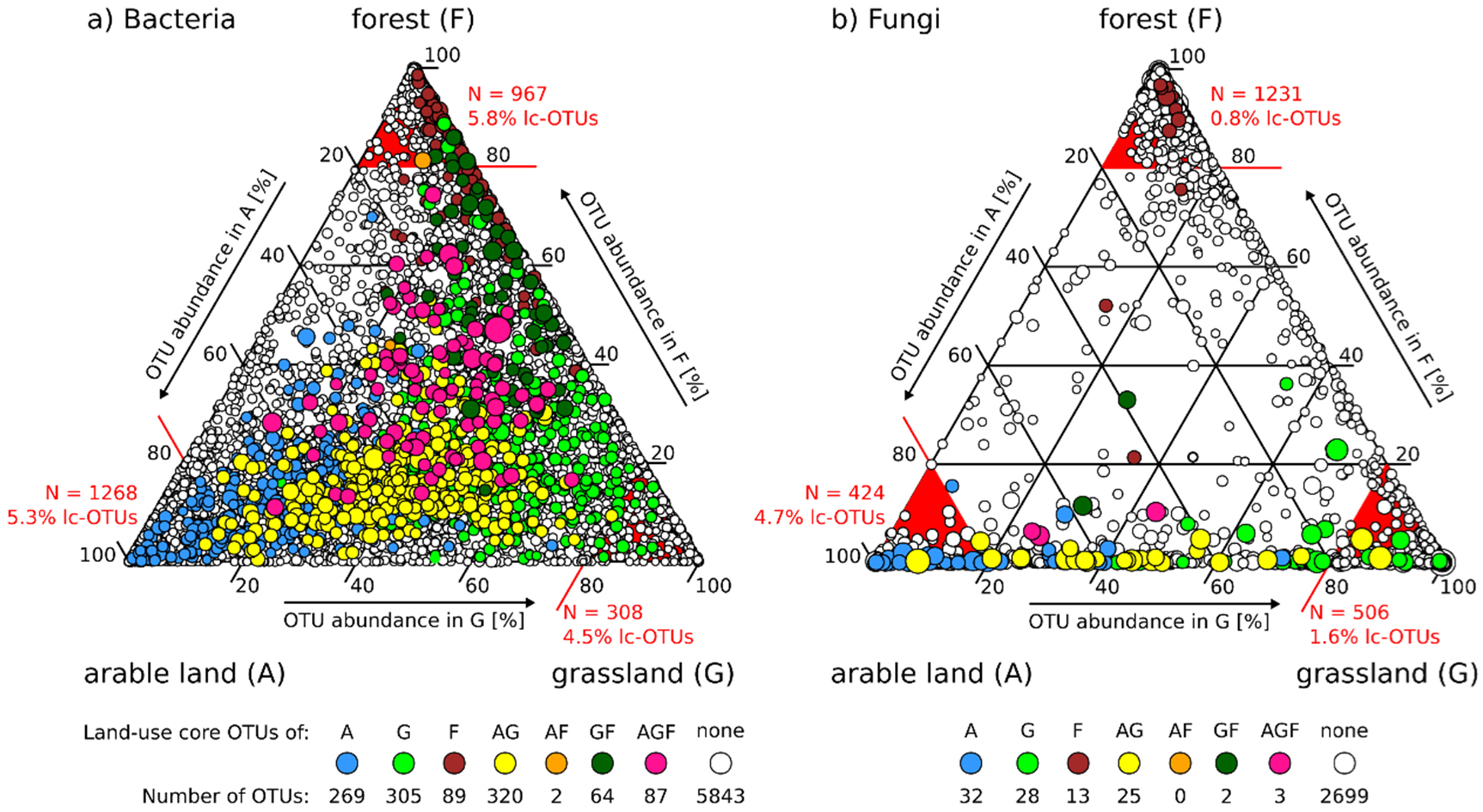
Ternary plots showing occurrences of the 6 979 bacterial (A) and 2 802 fungal (B) site core OTUs in the three land-use types and their combinations. Circles represent site core OTUs (sc-OTUs) and circle sizes indicate their relative abundance. Colored OTUs represent sc-OTUs, which are also land-use type core OTUs (lc-OTUs) from individual or combination of land-use types. White circles correspond to sc-OTUs, which are not part of land-use type core communities. The numbers of lc-OTUs of each land-use type or land-use type combinations are indicated below the ternary plots. The sc-OTUs were defined as OTUs occurring in 12 of 15 samples from a site and lc-OTUs as OTUs, which are sc-OTUs in 8 of 10 sites from a land-use type. Red lines and red triangles highlight the plot area, in which sc-OTUs occur which obtain at least 80% of their sequences from the respective single land-use type. The number of these sc-OTUs and the percent of lc-OTUs among these are indicated in red at the corners of the ternary plots.

### 3.5 Distribution of bacterial and fungal families among land-use types

For taxonomic characterization of core communities, we focused on the family level, since the classified OTUs can be more reliably assigned at this level and since the number of unclassified OTUs increased at lower taxonomic levels. For instance, 50.7% of the bacterial and 47.1% of the fungal OTUs were unclassified at the family-level, while these numbers were 78.0% for bacterial and 60.3% for fungal OTUs at the genus-level. In order to analyze associations of families to land-use types, we extracted sc-OTUs that were predominantly associated to a single or combinations of land-use types based on the ternary plot (Figure 4a). The ten most abundant families in each of the seven areas specified in the ternary plot, i.e., triangles A, G, F, AG, GF, AF, and AGF, were extracted. They covered in the selected areas 18.7% and 49.2% of the overall relative abundance of bacterial and fungal sc-OTUs, respectively, (Figure 4b, dark grey area) and resulted in a list of 39 bacterial and 38 fungal families (Figure 4c & 4d). Cluster analysis was used to group these families according to their distribution patterns in the land-use types, which yielded seven bacterial and five fungal clusters (Figure 4c & 4d). More homogenous representations of land-use types within the clusters were found for fungi as compared to bacteria. Most strikingly, fungal cluster V, which was composed of families such as Myxotrichaceae, Inocybaceae, and Russulaceae, occurred most strongly and almost exclusively in forest soils. Clusters predominantly associated to permanent grassland included only one bacterial family, the Ktedonobacteraceae (cluster IV, Figure 4c), but eight fungal families, e.g., Mortierellaceae and Chaetothyriaceae (cluster IV, Figure 4d). Within the clusters, also groupings with more resolved land-use type associations were revealed. For instance, within fungal cluster IV the fungal families Mortierellaceae Clavariaceae and Herpotrichiellaceae were all most abundant in permanent grassland but revealed a complex occurrence pattern in many land-use types, while the fungal family Chaetothyriaceae was exclusively detected in permanent grassland soils. Similarly, within fungal cluster III, which was mainly associated to arable land, some families such as Lasiosphaeriaceae and Nectriaceae were also prominently detected in the combination ‘arable land and permanent grassland’ while the Bulleribasidiaceae, as an exemption in cluster III, were more abundant in the combination ‘arable land and permanent grassland’ but comparably abundant in ‘arable land’. For bacteria, such clear clustering was less pronounced. Cluster VI exclusively associated to ‘arable land’ but for instance in cluster VII only eleven of the thirteen families were most abundant in forest soils. Within cluster VII, families such as Pedosphaeraceae or the candidate WD2101 soil group were also commonly detected in arable and permanent grassland soils. The strongest forest associations were observed for families Acidobacteriaceae Subgroup 1 as well as Acetobacteraceae, Methylacidiphilaceae, Acidothermaceae, and Micropepsaceae. Therefore, stronger associations to land-use types or their combinations were detected for fungi as compared to bacteria. This was further supported by the number of families with their highest abundance in a single land-use type (A, G, or F), which was lower for bacteria (20, Figure 4c) as compared to fungi (30, Figure 4d).

**Figure 4:**
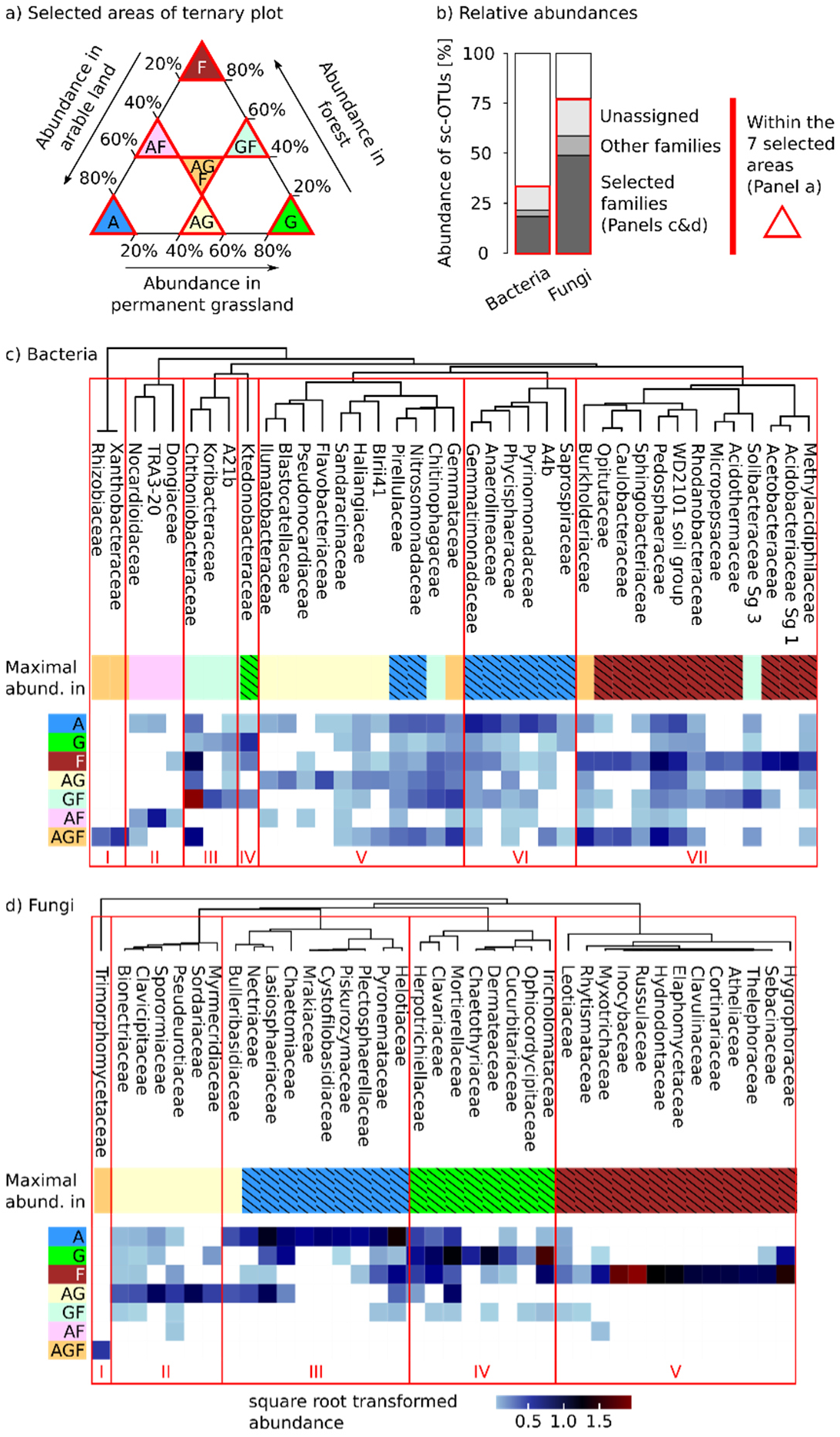
Distribution of most abundant bacterial and fungal families among land-use types. Based on the ternary plots (Figure 3) site-core OTUs (sc-OTUs) were selected from seven areas (a) corresponding to sc-OTUs with at least 80% of their abundance in a single land-use type (A, G, F), with at least 40% in each of two land-use types (AG, GF, AF), or with at least 20% in each land-use types (AGF). The proportions of relative abundances covered by the selected sc-OTUs, and their assignment at the family level, is shown in panel b). Panels c) and d) show the relative abundances of the ten most abundant bacterial (c) and fungal (d) families of each area of the ternary plots. Light blue indicates low, dark blue middle, and brown high relative abundances. White areas represent absences of families in an area of the ternary plot. The area in which a family has its highest abundance is indicated by the following color code (Maximal abund.): blue (A), green (G), brown (F), yellow (AG), light green (GF), pink (AF), and orange (AGF). Highest abundances in a single land-use type are indicated by black hatching. Dendrograms show clustering of normalized relative abundances of families in the land-use types and their combinations using average clustering (UPGMA). Red boxes highlight clusters of families with similar distributions among land-use types.

To detect families, which showed the strongest and most consistent associations to land-use types, we compared core and indicative OTUs. More specifically, we first selected OTUs, which were core and indicative of the same land-use type or land-use type combinations and aggregated these OTUs at the family-level. This yielded 304 bacterial (Table S7) and 58 fungal OTUs (Table S8). Then, we selected families, which included at least four (Bacteria) or two (Fungi) OTUs that were both core and indicative of the same land-uses (Table 5). This resulted in 16 bacterial and 9 fungal families (Table 5), which were also among the families described in Figure 4, with the exception of bacterial candidate groups SC-I-84 and AKYH767, as well as the fungal family Phaeosphaeriaceae. Two bacterial families, Anaerolineaceae and Pyrinomonadaceae included arable core and indicative OTUs and a single bacterial family, Acidobacteriaceae Subgroup 1, included only forest core and indicative OTUs. No bacterial family included only OTUs that were core and indicative of permanent grassland soils. Among fungi Chaetomiaceae and Myxotrichaceae included only OTUs that were core and indicative of a single land-use type, i.e., arable land and forest, respectively. No fungal family included exclusively OTUs that were core and indicative of permanent grassland soils. Furthermore, no bacterial and fungal OTUs were core and indicative of the combination ‘arable land and forest’ and only bacterial but no fungal families included OTUs that were core and indicative of ‘permanent grassland and forest’. The lack of such OTUs is consistent with the few sc-OTUs detected in the corresponding areas of the ternary plots (Figure 3), as well as with low similarities of bacterial and fungal communities among arable and forest sites, and equally low similarities among fungal communities of permanent grassland and forest sites (Figure 2).

**Table 5:** Number of OTUs, which were indicative (IndVal >0.8, p < 0.05) and core for the same land-use types from selected bacterial and fungal families. Families were selected if at least four (bacteria) or two (fungi) OTUs were indicative and core for the same land-use type or land-use type combination. All families are shown in Table S7 (Bacteria) and S8 (Fungi). Associations of families to land-use types are indicated according to Figure 4. Stars indicate families, which have the highest number of indicative and lc-OTUs and the highest abundance in the same land-use type or land-use type combination.

| Family | Indicative and lc-OTUs<sup>1)</sup> (all indicative OTUs) |  |  |  |  |  | Figure 4 |  |  |
| --- | --- | --- | --- | --- | --- | --- | --- | --- | --- |
|  | A | AG | G | AF | GF | F | Cluster | Main abund. |  |
| **Bacteria** |  |  |  |  |  |  |  |  |  |
| Chthoniobacteraceae | 1(1) | 4(12) | 0(1) | 0(0) | 2(5) | 3(4) | III | GF |  |
| Pirellulaceae | 4(5) | 1(8) | 0(0) | 0(0) | 0(2) | 0(0) | V | A | * |
| Chitinophagaceae | 4(10) | 3(12) | 2(2) | 0(0) | 1(4) | 1(3) | V | GF |  |
| Gemmatimonadaceae | 5(9) | 3(5) | 0(0) | 0(0) | 0(2) | 1(1) | VI | A | * |
| Anaerolineaceae | 4(4) | 0(3) | 0(0) | 0(0) | 0(1) | 0(0) | VI | A | * |
| Pyrinomonadaceae | 4(4) | 0(0) | 0(0) | 0(0) | 0(1) | 0(0) | VI | A | * |
| Burkholderiaceae | 1(2) | 4(7) | 0(0) | 0(0) | 0(1) | 3(4) | VII | AGF |  |
| Pedospaeraceae | 4(5) | 7(11) | 3(3) | 0(0) | 1(12) | 3(5) | VII | F |  |
| WD2101 soil group | 2(5) | 9(18) | 1(2) | 0(0) | 1(4) | 1(5) | VII | F |  |
| Acidobacteriaceae Sg 1 | 0(0) | 0(0) | 0(0) | 0(0) | 0(1) | 6(8) | VII | F | * |
| Acetobacteraceae | 0(0) | 0(0) | 0(0) | 0(0) | 2(2) | 4(6) | VII | F | * |
| Caulobacteraceae | 0(0) | 1(1) | 0(0) | 0(0) | 0(0) | 3(5) | VII | F | * |
| Acidothermaceae | 0(0) | 0(0) | 0(0) | 0(0) | 2(3) | 2(4) | VII | F | * |
| Solibacteraceae Sg 3 | 0(0) | 0(2) | 0(0) | 0(0) | 3(12) | 1(3) | VII | GF | * |
| AKYH767 | 0(0) | 4(4) | 0(0) | 0(0) | 0(0) | 0(0) |  |  |  |
| SC-I-84 | 1(1) | 2(4) | 2(2) | 0(0) | 0(3) | 0(0) |  |  |  |
| Others (incl. unclassified) | 47(75) | 53(195) | 12(21) | 0(4) | 28(110) | 17(41) |  |  |  |
| All | 86(128) | 105(287) | 23(29) | 0(4) | 42(162) | 48(89) |  |  |  |
| **Fungi** |  |  |  |  |  |  |  |  |  |
| Pseudeurotiaceae | 0(0) | 1(1) | 0(0) | 0(0) | 0(0) | 1(1) | II | AG | * |
| Lasiosphaeriaceae | 3(4) | 2(7) | 0(1) | 0(0) | 0(0) | 0(0) | III | A | * |
| Plectosphaerellaceae | 2(3) | 1(1) | 0(0) | 0(0) | 0(0) | 0(0) | III | A | * |
| Chaetomiaceae | 2(2) | 0(0) | 0(1) | 0(0) | 0(0) | 0(0) | III | A | * |
| Nectriaceae | 1(1) | 3(9) | 0(0) | 0(0) | 0(1) | 0(0) | III | A |  |
| Helotiaceae | 0(2) | 3(4) | 0(0) | 0(0) | 0(0) | 0(1) | III | A |  |
| Mortierellaceae | 0(0) | 2(5) | 1(1) | 0(0) | 0(0) | 1(1) | IV | G |  |
| Myxotrichaceae | 0(0) | 0(0) | 0(0) | 0(0) | 0(0) | 2(4) | V | F | * |
| Phaeosphaeriaceae | 1(1) | 1(1) | 0(0) | 0(0) | 0(0) | 0(0) |  |  |  |
| Others (incl. unclassified) | 7(35) | 12(53) | 5(13) | 0(2) | 0(0) | 7(20) |  |  |  |
| All | 16(47) | 25(79) | 6(16) | 0(2) | 0(1) | 11(26) |  |  |  |

<sup>1)</sup> indicative and lc-OTUs: OTUs, which are indicative and core of the same land-use type(s), note that OTUs cannot be indicative of all sites, i.e., the combination AGF;
<sup>2)</sup> Main abund.: Main abundance in arable land (A), permanent grassland (G), forest (F) or their combinations, according to Figure 4.

## 4. Discussion

### 4.1 Land-use-specificity of soil bacterial and fungal communities

Soil bacterial and fungal communities were surveyed during five years at thirty sites of the Swiss Soil Monitoring Network including three different land-use types, i.e., arable land, permanent grassland, and forest. This revealed communities that were highly specific to land-use types and sites, and which were stable over five years. A detailed analysis on the temporal stability of these communities has already been described (Gschwend *et al*. submitted). Here, we focused on the environmental drivers that shape this land-use- and site-specificity of soil bacterial and fungal communities, as well as on their taxonomic compositions.

Each land-use type was characterized by differences in the combinations of soil properties, management, and vegetation (Table S1). In arable soils, pH and bulk density were increased, while carbon contents were equal or lower than in permanent grassland and forest soils. Furthermore, management of arable soils included crop rotations, tillage (except one site), mineral and organic fertilization, as well as plant protection, which are known to influence soil bacterial and fungal communities (Hartmann *et al*. 2015, Peralta *et al*. 2018, Rivera-Becerril *et al*. 2017). Microbial biomass was significantly reduced in arable soils as compared to permanent grassland and forest soils (Table S1), which confirms earlier findings (Dequiedt *et al*. 2011). Bacterial communities in arable soils were characterized by families such as Anaerolineaceae, Pyrinomonadaceae, and Gemmatimonadaceae. Anaerolineaceae are widely distributed in soils, and particularly prevalent in low-oxygen environments, e.g., in compacted soils (Hartmann *et al*. 2014) or paddy fields (Jiao *et al*. 2019). As they may act as indicators for soil oxygen depletion (Gschwend *et al*. 2020), their high abundance in arable soils may be a sign of soil compaction in arable land due to common management practices with heavy machinery. Fungal communities in arable soils were for instance characterized by Lasiosphaeriaceae, Plectosphaerellaceae, Chaetomiaceae, and Mrakiaceae. With the exception of the basidiomycetous yeasts Mrakiaceae and Cystofilobasidiaceae (Liu *et al*. 2015), fungal families associated to arable soils also occurred in permanent grassland soils (Figure 4). For instance, Plectosphaerellaceae that include important soil-borne plant pathogens such as *Verticillium* (Giraldo and Crous 2019) had two lc-OTUs that were also indicative for arable land, as well as one that was indicative for ‘arable land and permanent grassland’ (Table 5). In these cases, OTUs assigned to the same family have distinct land-use type associations, which may for instance be driven by species-specific host plant preferences (Klosterman *et al*. 2009).

Permanent grassland soils were characterized by soil property values, which lay between those of arable and forest soils (Table S1). Their management included fertilization, mowing, and grazing, which may change soil bacterial and fungal community structures (Cui *et al*. 2020, Gilmullina *et al*. 2020, Kaiser *et al*. 2016). A single bacterial family, the Ktedonobacteraceae (phylum Chloroflexi) had their highest abundance in the permanent grassland section of the ternary plot, but also occurred in forest and less in arable soils (Figure 4c). Ktedonobacteraceae are aerobic, mycelium-forming bacteria and contain a single genus with one described species, i.e., *Ktedonobacter racemifer*, which was isolated from soil of a black locust forest in Italy (Cavaletti *et al*. 2006). Metabarcoding of bacterial communities from 2 173 soil samples across France revealed sequences assigned to *Ktedonobacter* in 80% of all samples, and attributed this genus to one of the dominant genera of soil bacteria (Karimi *et al*. 2018). Families that characterized fungal communities in permanent grasslands included for instance the grassland-specific Chaetothyriaceae (Figure 4d). Chaetothyriaceae include mainly epiphytic species living on plants (Quan *et al*. 2020) suggesting that their distribution may depend on host plants. However, in a survey of switchgrass-associated fungal communities, OTUs attributed to this family have also been detected associated to the switchgrass roots and adjacent soils, but not on plant leaves (Lee and Hawkes 2020), indicating that Chaetothyriaceae also include soil fungi.

Forest soils were characterized by relatively high contents of carbon, higher C/N-ratios, and lower soil pH as compared to the arable soils (Table S1). Bacterial families associated to forest soils included Acidobacteriaceae Subgroup 1, Acetobacteraceae, Acidothermaceae, as well as the more widely distributed WD2101 soil group, and Pedosphaeraceae (Figure 4c, Table 5). Acidobacteriaceae Subgroup 1 have been repeatedly reported to negatively correlate with soil pH (Kielak *et al*. 2016) and revealed increased abundances in soils with a pH below 6.5 (Jones *et al*. 2009). Acetobacteraceae have also been reported to strongly and negatively correlate with soil pH and to have higher abundances in forest as compared to grassland soils (Nacke *et al*. 2011). Therefore, soil pH, which is well known to be a major driver of soil bacterial communities (e.g. Karimi *et al*. 2018, Lauber *et al*. 2009), was the main factor determining forest associated soil bacterial taxa. Fungal communities in forest soils were mainly composed of ectomycorrhizal families such as Russulaceae, Inocybaceae, and Clavulinaceae, which is in agreement with previous findings (e.g. Frey *et al*. 2021). Thirteen fungal families were strongly associated to forest (Cluster V, Figure 4d), but only one of these, the Myxotrichaceae, included indicative OTUs of forest soils (Table S8). This is likely explained by the different forest ecosystems including deciduous, mixed and coniferous forests that have been sampled. As ectomycorrhizal fungi depend on their host tree species (Bahnmann *et al*. 2018), none of these families occurred at eight or more forest sites and were thus not generally indicative for forest soils. Myxotrichaceae included for instance *Oidiodendron* spp., which were repeatedly detected among the abundant soil fungi in metabarcoding surveys of Swiss forest soils (Frey *et al*. 2020, Hartmann *et al*. 2017), and which are common saprobes in acid soils but some of which also form ericoid mycorrhiza (Rice and Currah 2005). Therefore, their widespread and indicative distribution in various forest soil ecosystem may relate to a dependence on understory vegetation, or on the general preference for acidic soils.

### 4.2 Similarities of soil bacterial and fungal communities among land-use types

The similarities among soil bacterial communities from different land-use types were lowest for the combination of arable land and forest (Figure 2), which was also the only land-use type combination for which no bacterial lc-OTU was indicative (Table 5). Similarities between soil bacterial communities from arable and permanent grassland soils corresponded to values observed between permanent grassland and forest soils (Figure 2). This suggests that soil bacterial communities represented a sequential order following the soil properties and the land-use intensity from arable land, to permanent grassland and forest. For fungi, similarities from communities of permanent grassland and forest soils were equally low as among communities of arable and forest soils (Figure 2). Furthermore, no fungal OTUs was found that was indicative and land-use core for the combination ‘permanent grassland and forest’ or the combination ‘arable land and forest’ (Table 5). Therefore, soil fungal, unlike bacterial, communities revealed little overlap (Bray-Curtis < 0.10, Figure 2) between permanent grassland and forest soils. Considering dissimilarities among communities as proxies for the transfer of soil microorganisms among sites allows describing the structure of their metacommunities (Beck *et al*. 2019, Wisnoski and Lennon 2021). In this view, soil bacterial communities of arable, permanent grassland, and forest soils formed a single metacommunity, which was characterized by a continuous change from arable land, to permanent grassland and forest. Soil fungal communities, however, formed two metacommunities, one created by fungal communities of arable and permanent grassland soils and the other by fungal communities of forest soils.

The distinct structures of soil bacterial and fungal metacommunities can be explained by different factors influencing their community assembly. On the one hand, bacterial communities were more strongly structured by soil properties and climatic factors as compared to soil fungal communities (Table 4). On the other hand, soil fungal communities were more strongly structured by vegetation as compared to soil bacterial communities. For instance, acidophilic bacterial families predominantly occurred in forest soils (Figure 4c), while ectomycorrhizal fungal families dominated soil fungal communities in forest soils (Figure 4d). Confirming our results Frey *et al*. (2021) reported stronger effects of tree species on fungal as compared to bacterial community structures. Stronger vegetation effects on soil fungal as compared to bacterial communities were also revealed in the other land-use types, as crops had a stronger effect on soil fungal as compared to bacterial community structures (Table S3 & S4), which is in agreement with the findings of Ai *et al*. (2018). Stronger legacy effects of different grassland mixtures on soil fungal as compared to soil bacterial communities have been described in a grassland field experiment (Fox *et al*. 2020), which further supports the stronger impact of plants on soil fungal as compared to bacterial communities.

### 4.3 Potential use of sc-OTUs to provide a temporally stable, cultivation-independent reference list of dominant taxa

Site core OTUs accounted for 38.5% of bacterial and 33.1% of fungal OTUs, but covered 95.9% and 93.2% of relative abundance (Table 3). As sc-OTUs occurred in at least four of the five years, the large majority of retrieved sequences, could be attributed to temporally stable OTUs. Furthermore, these sc-OTUs not only were temporally stable but also included 95% of bacterial and 90% of fungal indicative OTUs (Figure S2) and were representative of the diversities of entire communities (Table S2). Therefore, sc-OTUs may be used to build a cultivation-independent, temporally stable reference set for the analysis of soil microbial diversity. Such reference sets are of particular interest for predictive modelling of soil bacterial and fungal diversity, and may also be used as reference values for long-term soil quality monitoring (Gschwend et al. submitted). Currently, long-term monitoring systems of soil biodiversity are largely lacking (Guerra *et al*. 2020, Leeuwen *et al*. 2017), which is particularly concerning given the ongoing environmental changes and the central role of soil biodiversity for global ecosystem processes. Finally, sc-OTUs provide support to establish lists of the most characteristic soil microorganisms, for which cultivation strategies or whole-genome sequencing are particularly valuable (Carini 2019). Currently, still too few dominating soil bacterial and fungal taxa have cultured representatives or available genome sequences, which would enable more detailed insight into their functions in the ecosystem (Delgado-Baquerizo *et al*. 2018, Egidi *et al*. 2019, Steen *et al*. 2019).

## 5. Conclusions

While microbial biomass and alpha-diversity measures at thirty long-term monitoring sites revealed only few differences among land-use types and sites, community compositions (Jaccard similarity) and structures (Bray-Curtis dissimilarity) yielded characteristic descriptors for each land-use type and site. Therefore, resolution obtained by metabarcoding were necessary to accurately describe soil bacterial and fungal communities. Temporally stable core OTUs accounted for 95.9% of bacterial and 93.2% of fungal sequences. These core OTUs were representative of entire communities and showed responses to distinct habitats. In total 4 184 indicative bacterial and 1 968 indicative fungal OTUs, of which 95% and 90% were also temporally stable core OTUs, were identified. These yield promising targets for the development of microbial indicators for robust soil quality analyses. Bacterial and fungal families were identified that revealed strong associations to one or more land-use types. In general, fungal families revealed stronger associations to land-use types, which may be explained by the stronger influences of vegetation on fungi as compared to bacteria, whereas bacteria were more strongly correlated with soil properties. Consequently, metacommunities of soil bacteria and fungi were differently structured. On the one hand, bacterial communities represented a sequential order following soil properties and land-use intensity from arable land, to permanent grassland and forest. On the other hand, fungal communities of forest sites showed only minor similarities to those from arable land and permanent grassland sites. The robustly assessed and temporally stable core OTUs may serve as references for future surveys of soil bacterial and fungal diversity. This may facilitate long-term soil quality monitoring by detecting disturbances of the characteristic habitat associated core communities, and it may also enable the development of predictive modelling for metabarcoding based soil quality analyses.

## 7. Acknowledgments

We thank Peter Schwab, Ramon Zimmermann, and further group members of the Swiss Soil Monitoring Network for soil sampling, Stephanie Pfister, Sonja Reinhard, and Beat Stierli for laboratory analyses, and Aaron Fox for valuable comments to the manuscript.

## Supplements

### Supplementary results

#### Sequence data overview

In total, 9 020 192 bacterial and 11 958 695 fungal high-quality sequences were obtained with at least 11 791 bacterial and 7 719 fungal sequences per sample and on average 20 045 (standard deviation: ± 3 705) bacterial and 26 575 (± 8 780) fungal sequences per sample. The average Good’s coverages were 0.92 (± 0.022) for bacteria and 0.98 (± 0.005) for fungi. Sequences were grouped into 18 140 bacterial OTUs (bOTU) and 8 477 fungal OTUs (fOTU) with an average of 2 714 (± 658) bOTUs and 562 (± 134) fOTUs per sample. Bacterial OTUs were assigned to 46 phyla, of which 31 occurred in core communities (Table S9) and fungal OTUs to 12 phyla of which 9 occurred in core communities (Table S10).

### Supplementary Figures and Tables

#### Supplementary figures

**Figure S1:**
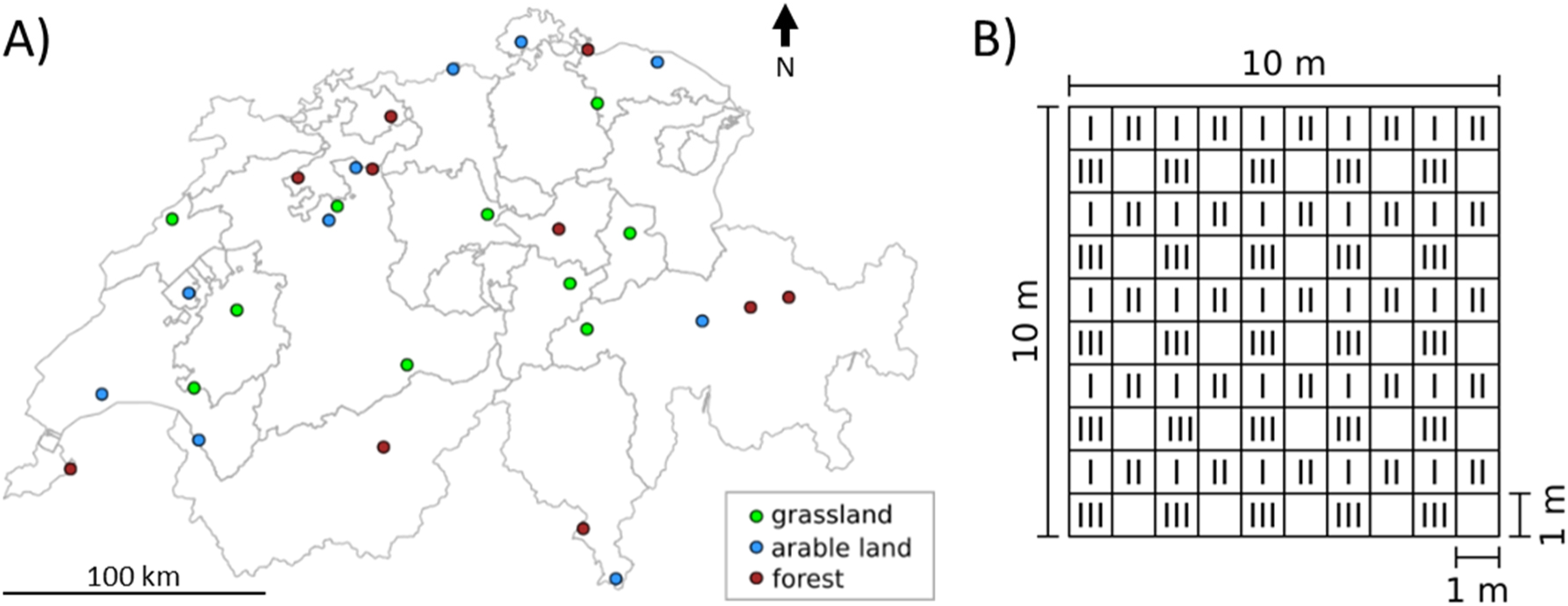
Map of Switzerland showing the thirty sampling sites and their land-use type (A). At each site three composite samples (I – III) were taken within a 10 m x 10 m area (B). Each composite sample was composed of 25 cores of 2.5 cm diameter and 20 cm depth. In each square meter marked with I to III, one core was randomly taken and mixed with other cores of the same number.

**Figure S2:**
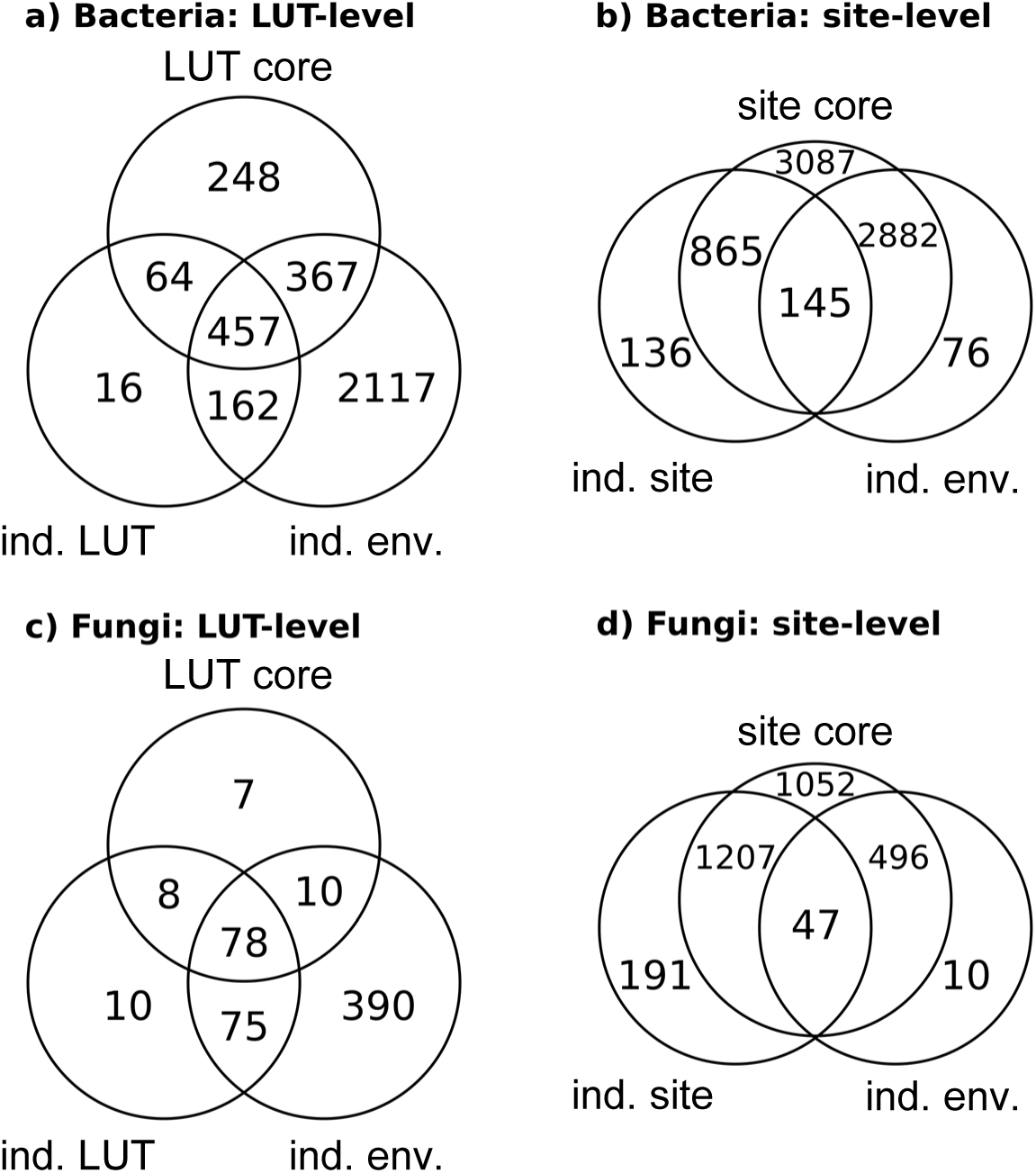
Venn diagramm depicting the OTUs that are shared between different OTU partitions. Bacterial (a, b) and fungal (c, d) OTU partitions are shown. Indicative and core OTUs were defined at two levels, i.e., land-use type (a, c) and site (b, d). Site core (sc) OTUs were defined as OTUs occurring in at least 80% of samples form a site, land-use type (LUT) cores (lc) as OTUs that were site cores of at least 80% of the sites from a land-use type. Indicative (ind.) OTUs were based on indicator species analysis (IndVal > 0.8), and environmental-factor-indicative-OTUs represent OTUs that revealed correlations to an environmental factor (|rho| > 0.4).

**Table S1:**
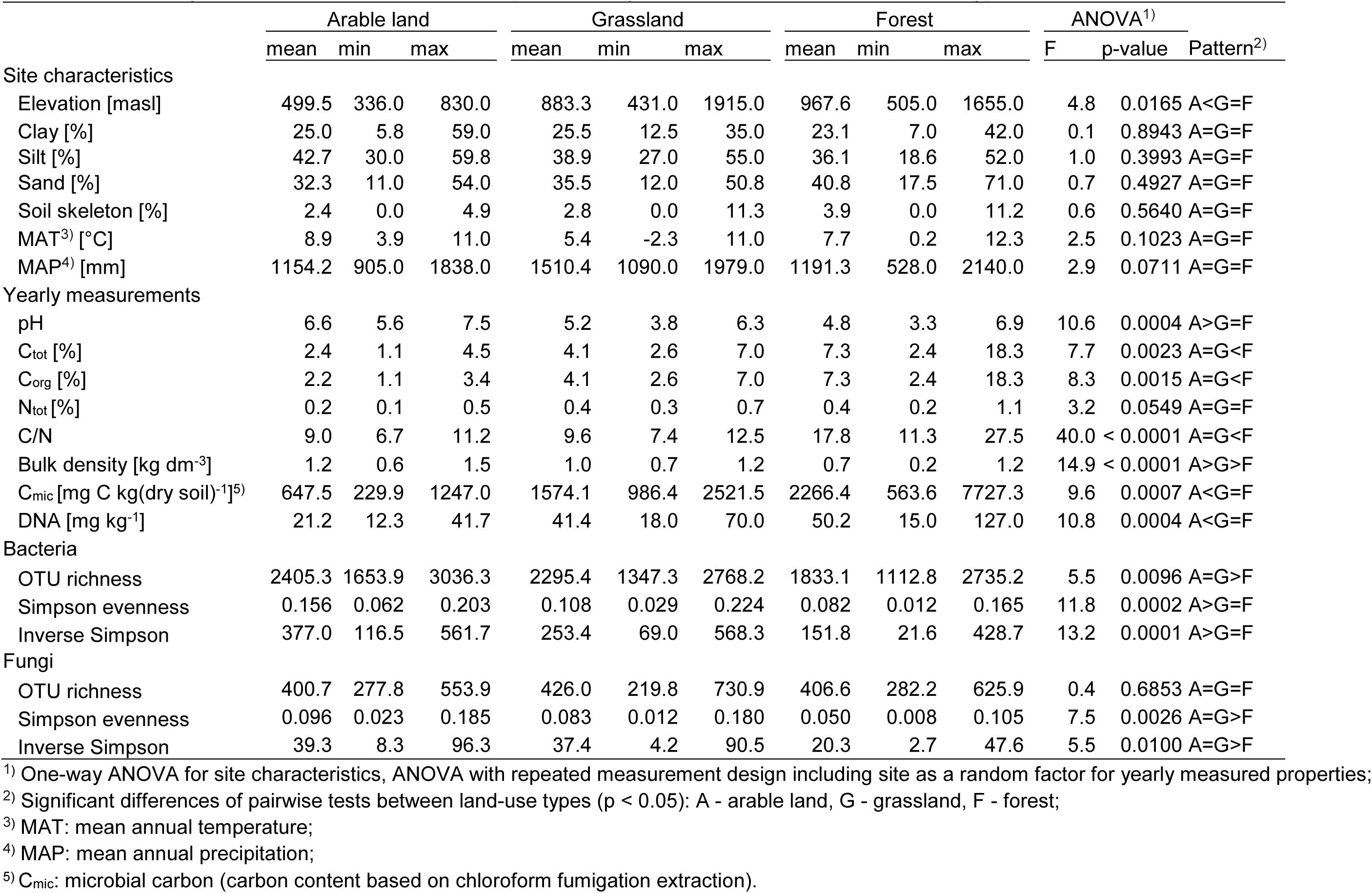
Summary of environmental factors and alpha diversity at the ten sites of each land-use type.

|  | Arable land |  |  | Grassland |  |  | Forest |  |  | ANOVA<sup>1)</sup> |  |  |
| --- | --- | --- | --- | --- | --- | --- | --- | --- | --- | --- | --- | --- |
|  | mean | min | max | mean | min | max | mean | min | max | F | p-value | Pattern<sup>2)</sup> |
| Site characteristics |  |  |  |  |  |  |  |  |  |  |  |  |
| Elevation [masl] | 499.5 | 336.0 | 830.0 | 883.3 | 431.0 | 1915.0 | 967.6 | 505.0 | 1655.0 | 4.8 | 0.0165 | A<G=F |
| Clay [%] | 25.0 | 5.8 | 59.0 | 25.5 | 12.5 | 35.0 | 23.1 | 7.0 | 42.0 | 0.1 | 0.8943 | A=G=F |
| Silt [%] | 42.7 | 30.0 | 59.8 | 38.9 | 27.0 | 55.0 | 36.1 | 18.6 | 52.0 | 1.0 | 0.3993 | A=G=F |
| Sand [%] | 32.3 | 11.0 | 54.0 | 35.5 | 12.0 | 50.8 | 40.8 | 17.5 | 71.0 | 0.7 | 0.4927 | A=G=F |
| Soil skeleton [%] | 2.4 | 0.0 | 4.9 | 2.8 | 0.0 | 11.3 | 3.9 | 0.0 | 11.2 | 0.6 | 0.5640 | A=G=F |
| MAT<sup>3)</sup> [°C] | 8.9 | 3.9 | 11.0 | 5.4 | -2.3 | 11.0 | 7.7 | 0.2 | 12.3 | 2.5 | 0.1023 | A=G=F |
| MAP<sup>4)</sup> [mm] | 1154.2 | 905.0 | 1838.0 | 1510.4 | 1090.0 | 1979.0 | 1191.3 | 528.0 | 2140.0 | 2.9 | 0.0711 | A=G=F |
| Yearly measurements |  |  |  |  |  |  |  |  |  |  |  |  |
| pH | 6.6 | 5.6 | 7.5 | 5.2 | 3.8 | 6.3 | 4.8 | 3.3 | 6.9 | 10.6 | 0.0004 | A>G=F |
| C<sub>tot</sub> [%] | 2.4 | 1.1 | 4.5 | 4.1 | 2.6 | 7.0 | 7.3 | 2.4 | 18.3 | 7.7 | 0.0023 | A=G<F |
| C<sub>org</sub> [%] | 2.2 | 1.1 | 3.4 | 4.1 | 2.6 | 7.0 | 7.3 | 2.4 | 18.3 | 8.3 | 0.0015 | A=G<F |
| N<sub>tot</sub> [%] | 0.2 | 0.1 | 0.5 | 0.4 | 0.3 | 0.7 | 0.4 | 0.2 | 1.1 | 3.2 | 0.0549 | A=G=F |
| C/N | 9.0 | 6.7 | 11.2 | 9.6 | 7.4 | 12.5 | 17.8 | 11.3 | 27.5 | 40.0 | < 0.0001 | A=G<F |
| Bulk density [kg dm<sup>-3</sup>] | 1.2 | 0.6 | 1.5 | 1.0 | 0.7 | 1.2 | 0.7 | 0.2 | 1.2 | 14.9 | < 0.0001 | A>G>F |
| C<sub>mic</sub> [mg C kg(dry soil)<sup>-1</sup>]<sup>5)</sup> | 647.5 | 229.9 | 1247.0 | 1574.1 | 986.4 | 2521.5 | 2266.4 | 563.6 | 7727.3 | 9.6 | 0.0007 | A<G=F |
| DNA [mg kg<sup>-1</sup>] | 21.2 | 12.3 | 41.7 | 41.4 | 18.0 | 70.0 | 50.2 | 15.0 | 127.0 | 10.8 | 0.0004 | A<G=F |
| Bacteria |  |  |  |  |  |  |  |  |  |  |  |  |
| OTU richness | 2405.3 | 1653.9 | 3036.3 | 2295.4 | 1347.3 | 2768.2 | 1833.1 | 1112.8 | 2735.2 | 5.5 | 0.0096 | A=G>F |
| Simpson evenness | 0.156 | 0.062 | 0.203 | 0.108 | 0.029 | 0.224 | 0.082 | 0.012 | 0.165 | 11.8 | 0.0002 | A>G=F |
| Inverse Simpson | 377.0 | 116.5 | 561.7 | 253.4 | 69.0 | 568.3 | 151.8 | 21.6 | 428.7 | 13.2 | 0.0001 | A>G=F |
| Fungi |  |  |  |  |  |  |  |  |  |  |  |  |
| OTU richness | 400.7 | 277.8 | 553.9 | 426.0 | 219.8 | 730.9 | 406.6 | 282.2 | 625.9 | 0.4 | 0.6853 | A=G=F |
| Simpson evenness | 0.096 | 0.023 | 0.185 | 0.083 | 0.012 | 0.180 | 0.050 | 0.008 | 0.105 | 7.5 | 0.0026 | A=G>F |
| Inverse Simpson | 39.3 | 8.3 | 96.3 | 37.4 | 4.2 | 90.5 | 20.3 | 2.7 | 47.6 | 5.5 | 0.0100 | A=G>F |

<sup>1)</sup> One-way ANOVA for site characteristics, ANOVA with repeated measurement design including site as a random factor for yearly measured properties;904 <sup>2)</sup> Significant differences of pairwise tests between land-use types (p < 0.05): A - arable land, G - grassland, F - forest;905 <sup>3)</sup> MAT: mean annual temperature;906 <sup>4)</sup> MAP: mean annual precipitation;907 <sup>5)</sup> C<sub>mic</sub>: microbial carbon (carbon content based on chloroform fumigation extraction).

**Table S2:** Spearman correlations of entire communities with those composed of site-core OTUs (sc-OTUs). OTUs being detected in at least 12 of the 15 samples from a site were classified as sc-OTUs.

|  | Bacteria |  | Fungi |  |
| --- | --- | --- | --- | --- |
|  | rho | p-value | rho | p-value |
| Alpha-diversity |  |  |  |  |
| OTU richness | 0.99 | < 2.2e-16 | 0.97 | < 2.2e-16 |
| Simpson evenness | 1.00 | < 2.2e-16 | 0.98 | < 2.2e-16 |
| Inverse Simpson | 1.00 | < 2.2e-16 | 0.98 | < 2.2e-16 |
| Beta diversity |  |  |  |  |
| Jaccard | 1.00 | 0.0001 | 1.00 | 0.0001 |
| Bray-Curtis | 1.00 | 0.0001 | 1.00 | 0.0001 |

**Table S3:**
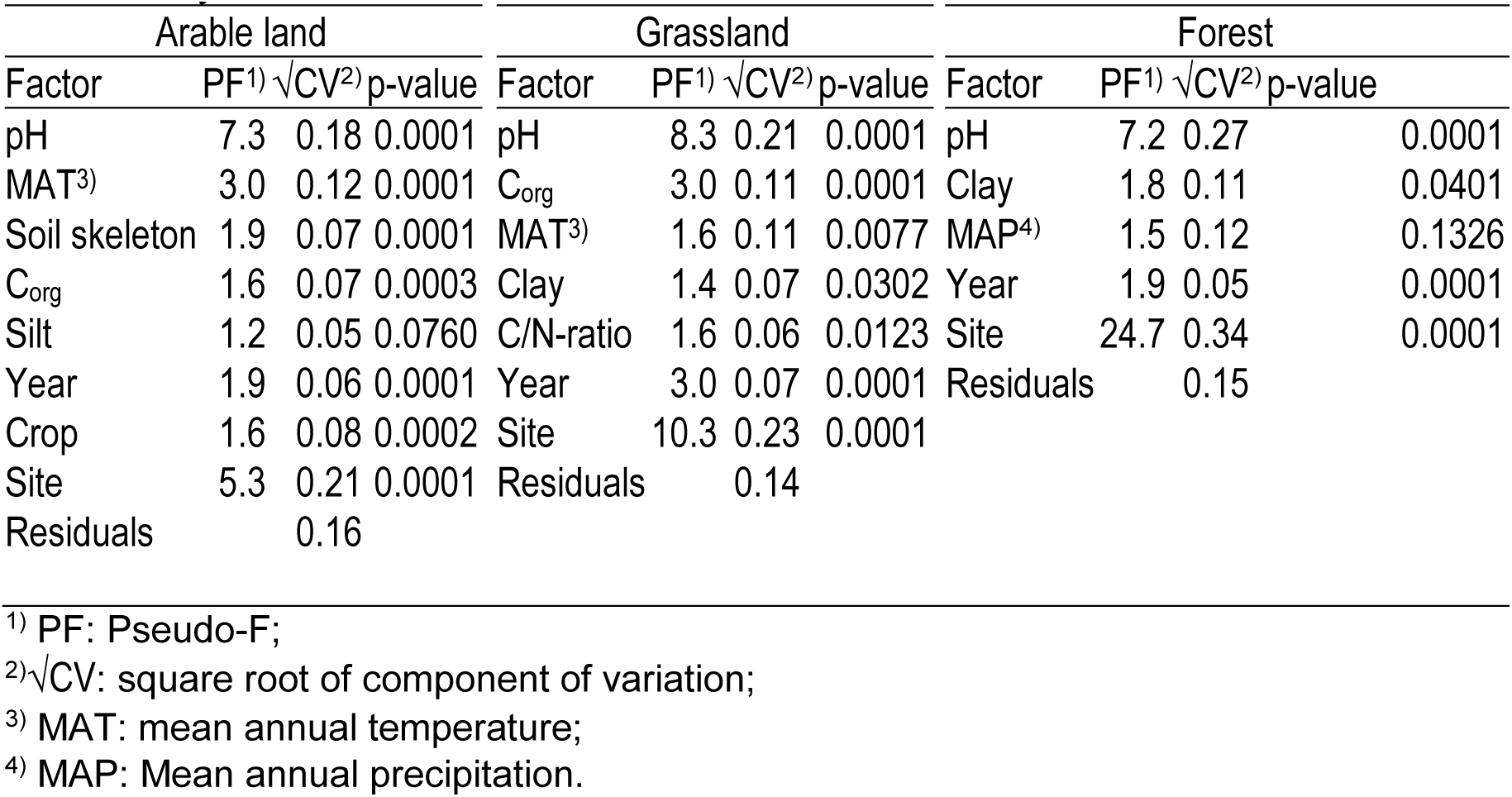
Environmental influences on bacterial communities in three land-use types as assessed by nested PERMANOVA. Model selection used AICc as selection criteria.

**Table S4:** Environmental influences on fungal communities in three land-use types as assessed by nested PERMANOVA. Model selection used AICc as selection criteria.

| Arable land |  |  |  | Grassland |  |  |  | Forest |  |  |  |
| --- | --- | --- | --- | --- | --- | --- | --- | --- | --- | --- | --- |
| Factor | PF<sup>1)</sup> | √CV<sup>2)</sup> | p-value | Factor | PF<sup>1)</sup> | √CV | p-value | Factor | PF<sup>1)</sup> | √CV<sup>2)</sup> | p-value |
| pH | 1.7 | 0.11 | 0.0050 | pH | 4.5 | 0.23 | 0.0001 | pH | 1.4 | 0.13 | 0.2000 |
| Year | 1.4 | 0.08 | 0.0025 | Altitude | 2.3 | 0.20 | 0.0009 | Site | 13.6 | 0.59 | 0.0001 |
| Crop | 1.5 | 0.15 | 0.0007 | Clay | 1.5 | 0.09 | 0.0553 | Residuals |  | 0.36 |  |
| Site | 6.6 | 0.38 | 0.0001 | Soil skeleton | 1.3 | 0.09 | 0.1192 |  |  |  |  |
| Residuals |  | 0.31 |  | C<sub>org</sub> | 1.6 | 0.18 | 0.0413 |  |  |  |  |
|  |  |  |  | Year | 2.3 | 0.09 | 0.0001 |  |  |  |  |
|  |  |  |  | Site | 10.2 | 0.38 | 0.0001 |  |  |  |  |
|  |  |  |  | Residuals |  | 0.23 |  |  |  |  |  |

<sup>1)</sup> PF: Pseudo-F;
<sup>2)</sup> √CV: square root of component of variation.

**Table S5:**
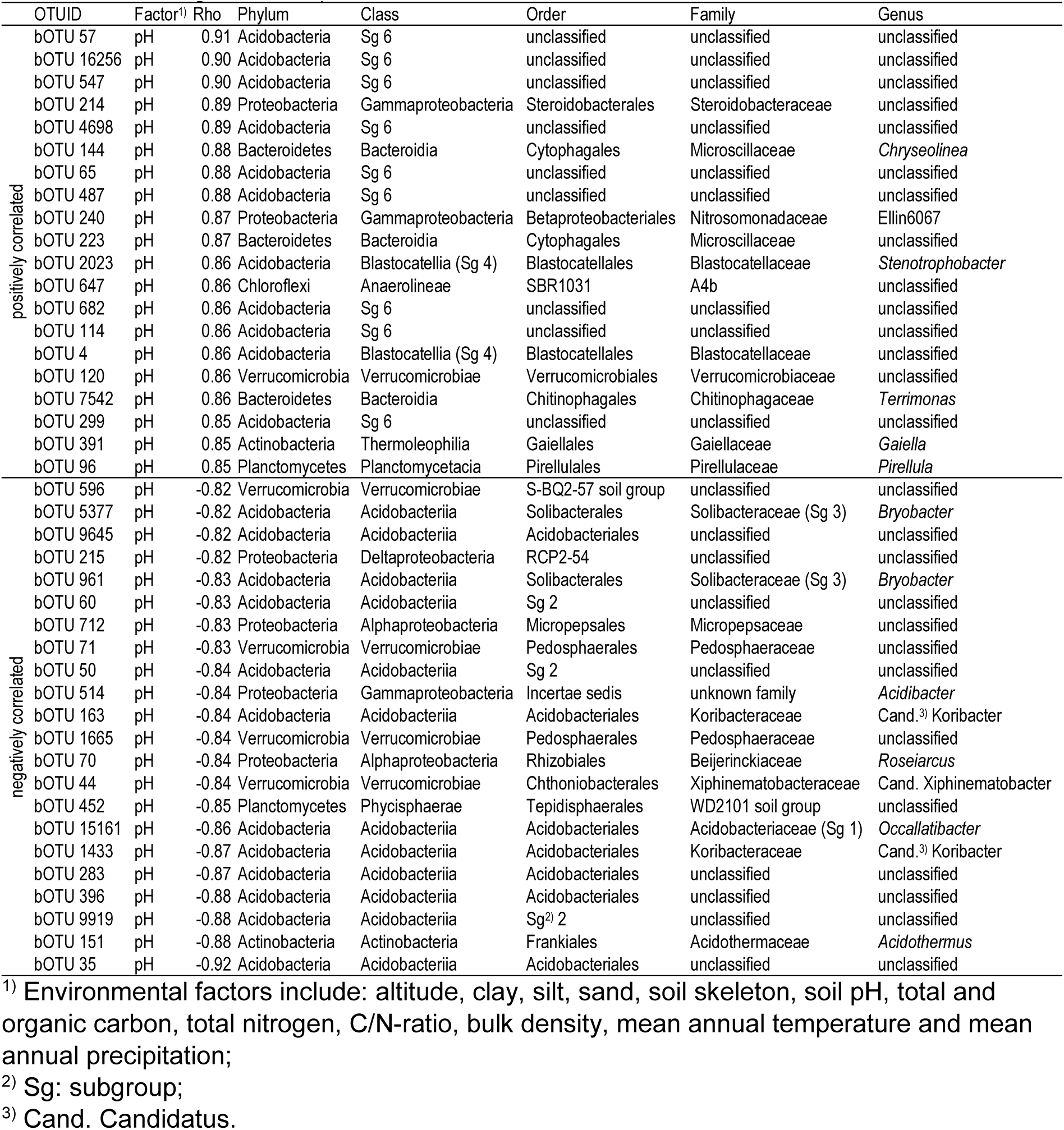
Bacterial OTUs with the strongest negative and positive correlations to an environmental factor along with their taxonomic assignment. The 20 strongest correlations are shown for negative and positive correlations.

**Table S6:**
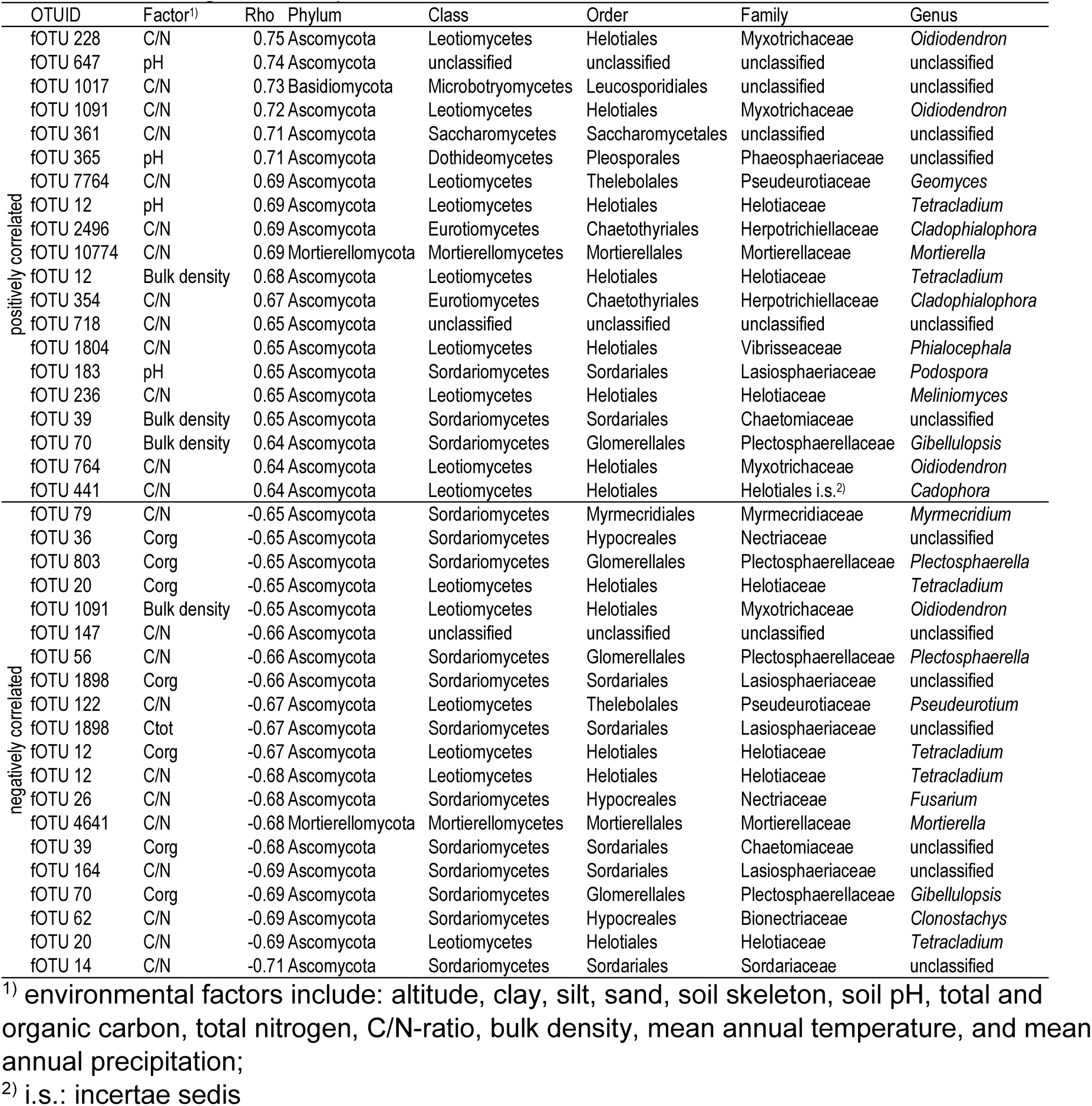
Fungal OTUs with the strongest negative and positive correlations to an environmental factor along with their taxonomic assignment. The 20 strongest correlations are shown for negative and positive correlations.

**Table S7:**
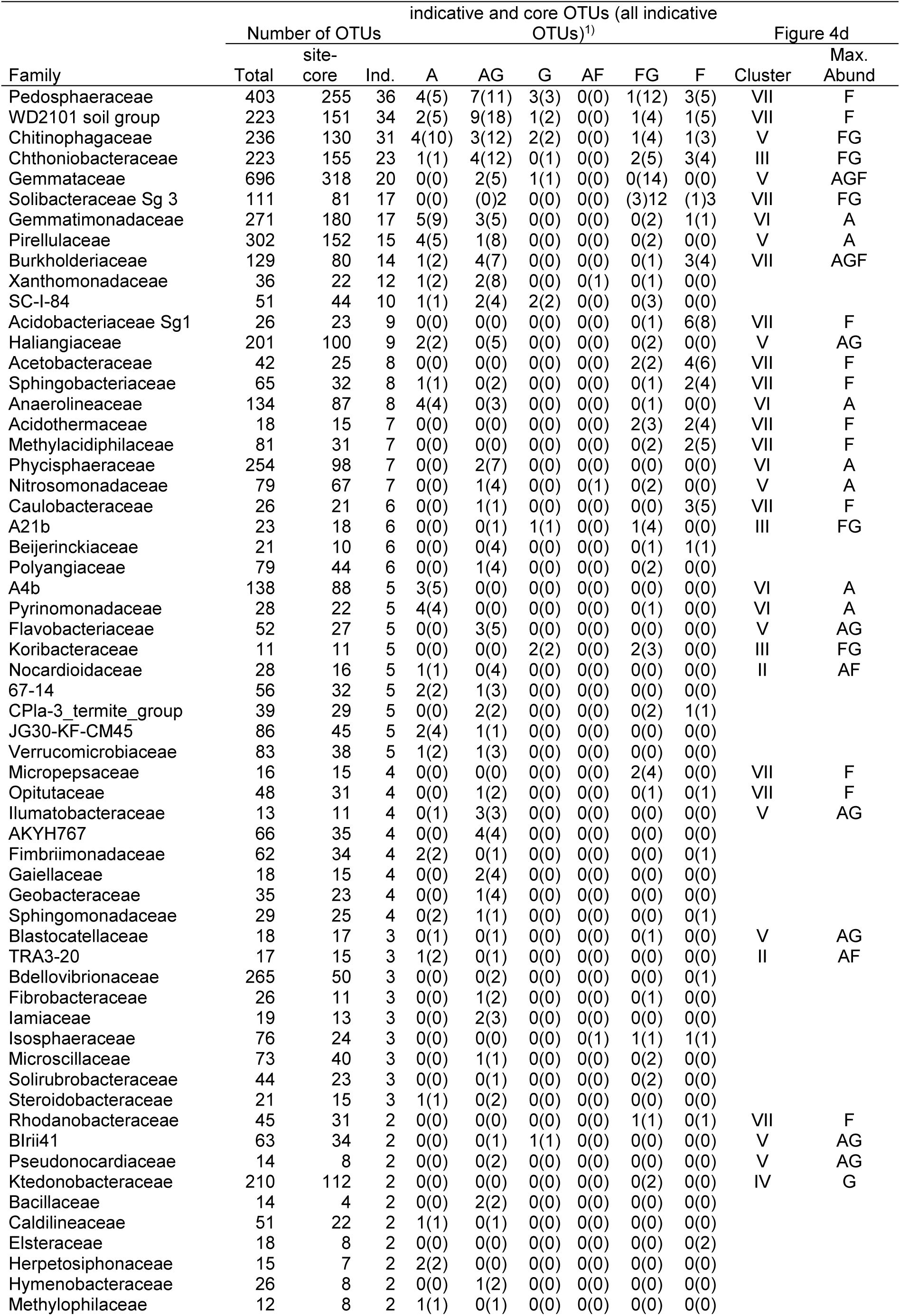
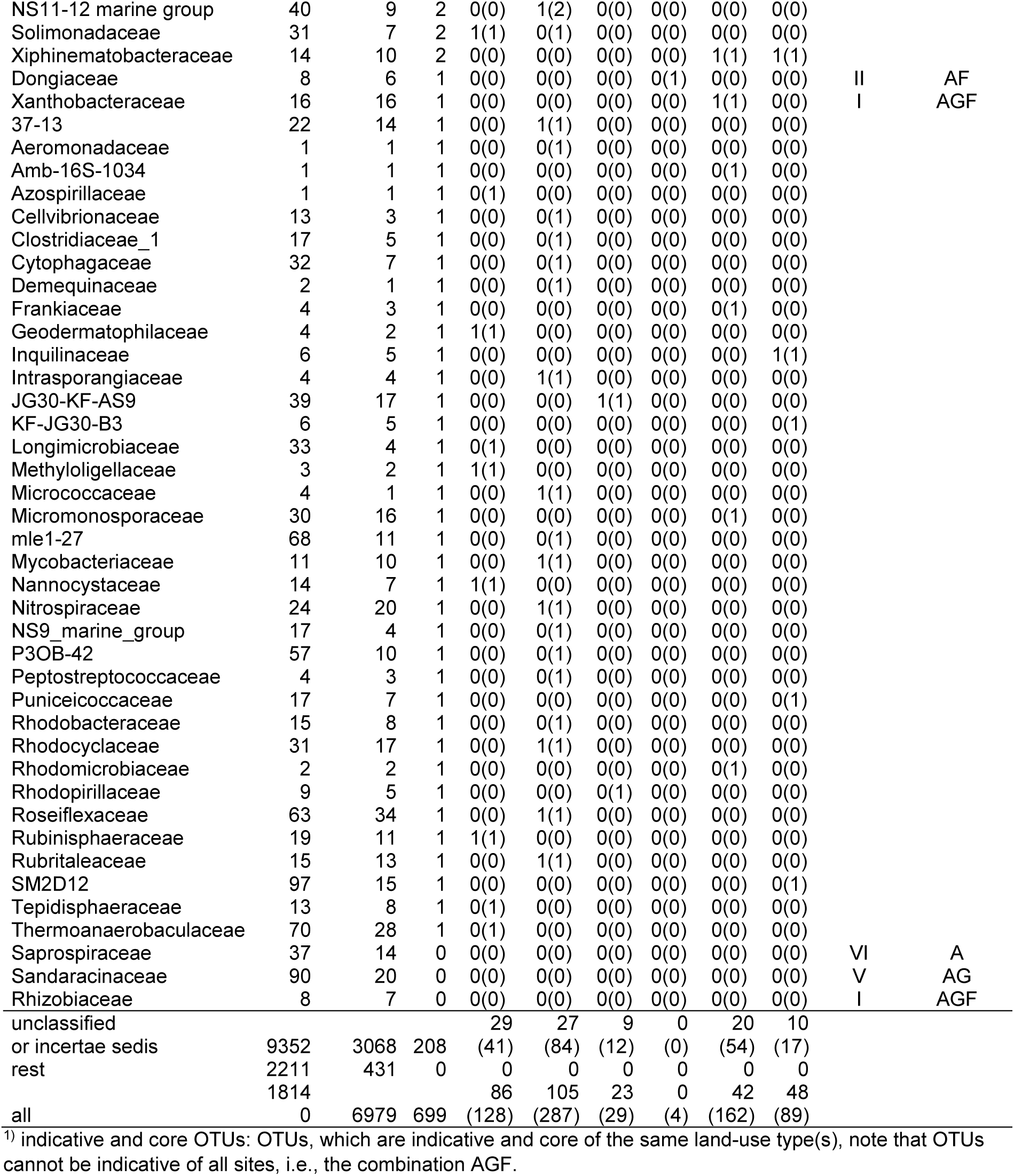
Number of land-use-indicative bacterial OTUs grouped by family. All families containing at least one indicative OTU, as well as families represented in Figure 4c are shown.

**Table S8:** Number of land-use-indicative fungal OTUs grouped by family. All families containing at least one indicative OTU, as well as families represented in Figure 4d are shown.

| Family | Number of OTUs |  |  | indicative and core OTUs (all indicative OTUs)<sup>1)</sup> |  |  |  |  |  | Figure 4d |  |
| --- | --- | --- | --- | --- | --- | --- | --- | --- | --- | --- | --- |
|  | Total | site-core | indicative | A | AG | G | AF | FG | F | Cluster | Max. Abund. |
| Lasiosphaeriaceae | 58 | 41 | 12 | 3(4) | 2(7) | 0(1) | 0(0) | 0(0) | 0(0) | III | A |
| Nectriaceae | 52 | 27 | 11 | 1(1) | 3(9) | 0(0) | 0(0) | 0(1) | 0(0) | III | A |
| Herpotrichiellaceae | 143 | 67 | 7 | 0(2) | 0(1) | 0(0) | 0(0) | 0(0) | 1(4) | IV | G |
| Mortierellaceae | 129 | 58 | 7 | 0(0) | 2(5) | 1(1) | 0(0) | 0(0) | 1(1) | IV | G |
| Helotiaceae | 120 | 44 | 7 | 0(2) | 3(4) | 0(0) | 0(0) | 0(0) | 0(1) | III | A |
| Hypocreaceae | 25 | 11 | 6 | 0(2) | 0(2) | 0(0) | 0(2) | 0(0) | 0(0) |  |  |
| Myxotrichaceae | 40 | 18 | 4 | 0(0) | 0(0) | 0(0) | 0(0) | 0(0) | 2(4) | V | F |
| Plectosphaerellaceae | 23 | 8 | 4 | 2(3) | 1(1) | 0(0) | 0(0) | 0(0) | 0(0) | III | A |
| Hyaloscyphaceae | 85 | 40 | 4 | 0(0) | 0(2) | 0(1) | 0(0) | 0(0) | 0(1) |  |  |
| Cucurbitariaceae | 14 | 9 | 3 | 0(0) | 0(2) | 1(1) | 0(0) | 0(0) | 0(0) | IV | G |
| Bulleribasidiaceae | 26 | 7 | 3 | 0(1) | 1(2) | 0(0) | 0(0) | 0(0) | 0(0) | III | AG |
| Chaetomiaceae | 39 | 20 | 3 | 2(2) | 0(0) | 0(1) | 0(0) | 0(0) | 0(0) | III | A |
| Mrakiaceae | 7 | 4 | 3 | 0(3) | 0(0) | 0(0) | 0(0) | 0(0) | 0(0) | III | A |
| Dermateaceae | 25 | 10 | 2 | 0(0) | 1(1) | 0(0) | 0(0) | 0(0) | 0(1) | IV | G |
| Tricholomataceae | 103 | 37 | 2 | 0(0) | 0(0) | 0(1) | 0(0) | 0(0) | 1(1) | IV | G |
| Clavicipitaceae | 35 | 19 | 2 | 0(0) | 0(2) | 0(0) | 0(0) | 0(0) | 0(0) | II | AG |
| Pseudeurotiaceae | 15 | 11 | 2 | 0(0) | 1(1) | 0(0) | 0(0) | 0(0) | 1(1) | II | AG |
| Chaetosphaeriaceae | 36 | 16 | 2 | 0(0) | 0(1) | 0(1) | 0(0) | 0(0) | 0(0) |  |  |
| Gloniaceae | 11 | 10 | 2 | 0(0) | 0(0) | 0(0) | 0(0) | 0(0) | 1(2) |  |  |
| Leucosporidiaceae | 7 | 3 | 2 | 0(0) | 0(2) | 0(0) | 0(0) | 0(0) | 0(0) |  |  |
| Phaeosphaeriaceae | 53 | 13 | 2 | 1(1) | 1(1) | 0(0) | 0(0) | 0(0) | 0(0) |  |  |
| Leotiaceae | 43 | 24 | 1 | 0(0) | 0(0) | 0(1) | 0(0) | 0(0) | 0(0) | V | F |
| Chaetothyriaceae | 4 | 2 | 1 | 0(0) | 0(0) | 0(1) | 0(0) | 0(0) | 0(0) | IV | G |
| Piskurozymaceae | 11 | 3 | 1 | 0(0) | 0(1) | 0(0) | 0(0) | 0(0) | 0(0) | III | A |
| Pyronemataceae | 92 | 38 | 1 | 0(0) | 0(1) | 0(0) | 0(0) | 0(0) | 0(0) | III | A |
| Bionectriaceae | 9 | 7 | 1 | 0(0) | 1(1) | 0(0) | 0(0) | 0(0) | 0(0) | II | AG |
| Myrmecridiaceae | 8 | 2 | 1 | 0(0) | 1(1) | 0(0) | 0(0) | 0(0) | 0(0) | II | AG |
| Sordariaceae | 4 | 1 | 1 | 0(0) | 1(1) | 0(0) | 0(0) | 0(0) | 0(0) | II | AG |
| Sporormiaceae | 23 | 5 | 1 | 0(0) | 1(1) | 0(0) | 0(0) | 0(0) | 0(0) | II | AG |
| Ascobolaceae | 13 | 7 | 1 | 0(0) | 0(1) | 0(0) | 0(0) | 0(0) | 0(0) |  |  |
| Aspergillaceae | 51 | 22 | 1 | 0(0) | 0(0) | 0(0) | 0(0) | 0(0) | 0(1) |  |  |
| Cystofilobasidiaceae | 5 | 2 | 1 | 0(1) | 0(0) | 0(0) | 0(0) | 0(0) | 0(0) | III | A |
| Didymellaceae | 7 | 2 | 1 | 0(0) | 1(1) | 0(0) | 0(0) | 0(0) | 0(0) |  |  |
| Didymosphaeriaceae | 6 | 3 | 1 | 0(0) | 0(1) | 0(0) | 0(0) | 0(0) | 0(0) |  |  |
| Filobasidiaceae | 10 | 1 | 1 | 0(1) | 0(0) | 0(0) | 0(0) | 0(0) | 0(0) |  |  |
| Hyponectriaceae | 4 | 2 | 1 | 0(0) | 0(1) | 0(0) | 0(0) | 0(0) | 0(0) |  |  |
| Leptosphaeriaceae | 19 | 5 | 1 | 0(1) | 0(0) | 0(0) | 0(0) | 0(0) | 0(0) |  |  |
| Magnaporthaceae | 17 | 5 | 1 | 0(0) | 0(1) | 0(0) | 0(0) | 0(0) | 0(0) |  |  |
| Melanommataceae | 5 | 2 | 1 | 0(0) | 0(1) | 0(0) | 0(0) | 0(0) | 0(0) |  |  |
| Microdochiaceae | 5 | 3 | 1 | 0(1) | 0(0) | 0(0) | 0(0) | 0(0) | 0(0) |  |  |
| Minutisphaeraceae | 6 | 2 | 1 | 0(0) | 0(0) | 1(1) | 0(0) | 0(0) | 0(0) |  |  |
| Mucoraceae | 21 | 9 | 1 | 0(0) | 0(1) | 0(0) | 0(0) | 0(0) | 0(0) |  |  |
| Periconiaceae | 6 | 2 | 1 | 1(1) | 0(0) | 0(0) | 0(0) | 0(0) | 0(0) |  |  |
| Pleosporaceae | 20 | 5 | 1 | 0(1) | 0(0) | 0(0) | 0(0) | 0(0) | 0(0) |  |  |
| Thyridariaceae | 2 | 1 | 1 | 0(0) | 0(1) | 0(0) | 0(0) | 0(0) | 0(0) |  |  |
| Torulaceae | 6 | 1 | 1 | 0(1) | 0(0) | 0(0) | 0(0) | 0(0) | 0(0) |  |  |
| Trichocomaceae | 18 | 9 | 1 | 0(1) | 0(0) | 0(0) | 0(0) | 0(0) | 0(0) |  |  |
| Trichosporonaceae | 7 | 3 | 1 | 0(0) | 1(1) | 0(0) | 0(0) | 0(0) | 0(0) |  |  |
| Venturiaceae | 38 | 22 | 1 | 0(0) | 0(0) | 0(0) | 0(0) | 0(0) | 1(1) |  |  |
| Vibrisseaceae | 10 | 5 | 1 | 0(0) | 0(0) | 0(0) | 0(0) | 0(0) | 0(1) |  |  |
| Atheliaceae | 42 | 26 | 0 | 0(0) | 0(0) | 0(0) | 0(0) | 0(0) | 0(0) | V | F |
| Clavulinaceae | 27 | 16 | 0 | 0(0) | 0(0) | 0(0) | 0(0) | 0(0) | 0(0) | V | F |
| Cortinariaceae | 129 | 48 | 0 | 0(0) | 0(0) | 0(0) | 0(0) | 0(0) | 0(0) | V | F |
| Elaphomycetaceae | 9 | 6 | 0 | 0(0) | 0(0) | 0(0) | 0(0) | 0(0) | 0(0) | V | F |
| Hydnodontaceae | 35 | 13 | 0 | 0(0) | 0(0) | 0(0) | 0(0) | 0(0) | 0(0) | V | F |
| Hygrophoraceae | 44 | 23 | 0 | 0(0) | 0(0) | 0(0) | 0(0) | 0(0) | 0(0) | V | F |
| Inocybaceae | 122 | 69 | 0 | 0(0) | 0(0) | 0(0) | 0(0) | 0(0) | 0(0) | V | F |
| Rhytismataceae | 20 | 6 | 0 | 0(0) | 0(0) | 0(0) | 0(0) | 0(0) | 0(0) | V | F |
| Russulaceae | 85 | 60 | 0 | 0(0) | 0(0) | 0(0) | 0(0) | 0(0) | 0(0) | V | F |
| Sebacinaceae | 64 | 43 | 0 | 0(0) | 0(0) | 0(0) | 0(0) | 0(0) | 0(0) | V | F |
| Thelephoraceae | 136 | 70 | 0 | 0(0) | 0(0) | 0(0) | 0(0) | 0(0) | 0(0) | V | F |
| Clavariaceae | 166 | 71 | 0 | 0(0) | 0(0) | 0(0) | 0(0) | 0(0) | 0(0) | IV | G |
| Ophiocordycipitaceae | 20 | 7 | 0 | 0(0) | 0(0) | 0(0) | 0(0) | 0(0) | 0(0) | IV | G |
| Trimorphomycetaceae | 5 | 2 | 0 | 0(0) | 0(0) | 0(0) | 0(0) | 0(0) | 0(0) | I | AGF |
| unclassified or incertae |  |  |  |  |  |  |  |  |  |  |  |
| sedis | 4256 | 1187 | 52 | 6(18) | 4(21) | 3(6) | 0(0) | 0(0) | 3(7) |  |  |
| Rest | 1801 | 487 | 0 | 0 | 0 | 0 | 0 | 0 | 0 |  |  |
| All | 8477 | 2802 | 171 | 16(47) | 25(79) | 6(16) | 0(2) | 0(1) | 11(26) |  |  |

<sup>1)</sup> indicative and core OTUs: OTUs, which are indicative and core of the same land-use type(s), note that OTUs cannot be indicative of all sites, i.e., the combination AGF.

**Table S9:**
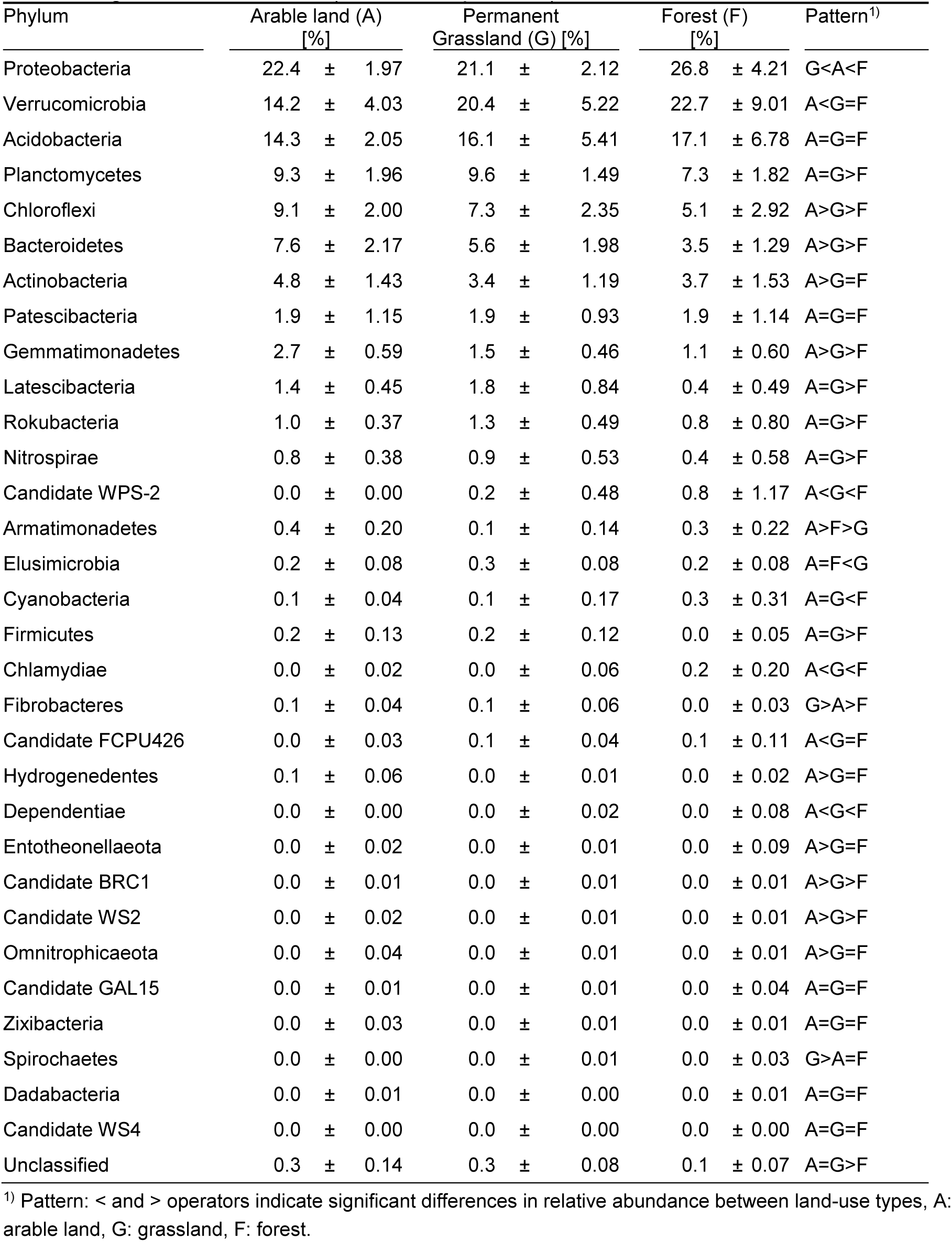
Relative abundances of bacterial phyla in arable land, permanent grassland, and forest. Mean values (± standard deviations) of each land-use type are shown. Different letters indicate significant differences (Dunn Test, p < 0.05).

**Table S10.**
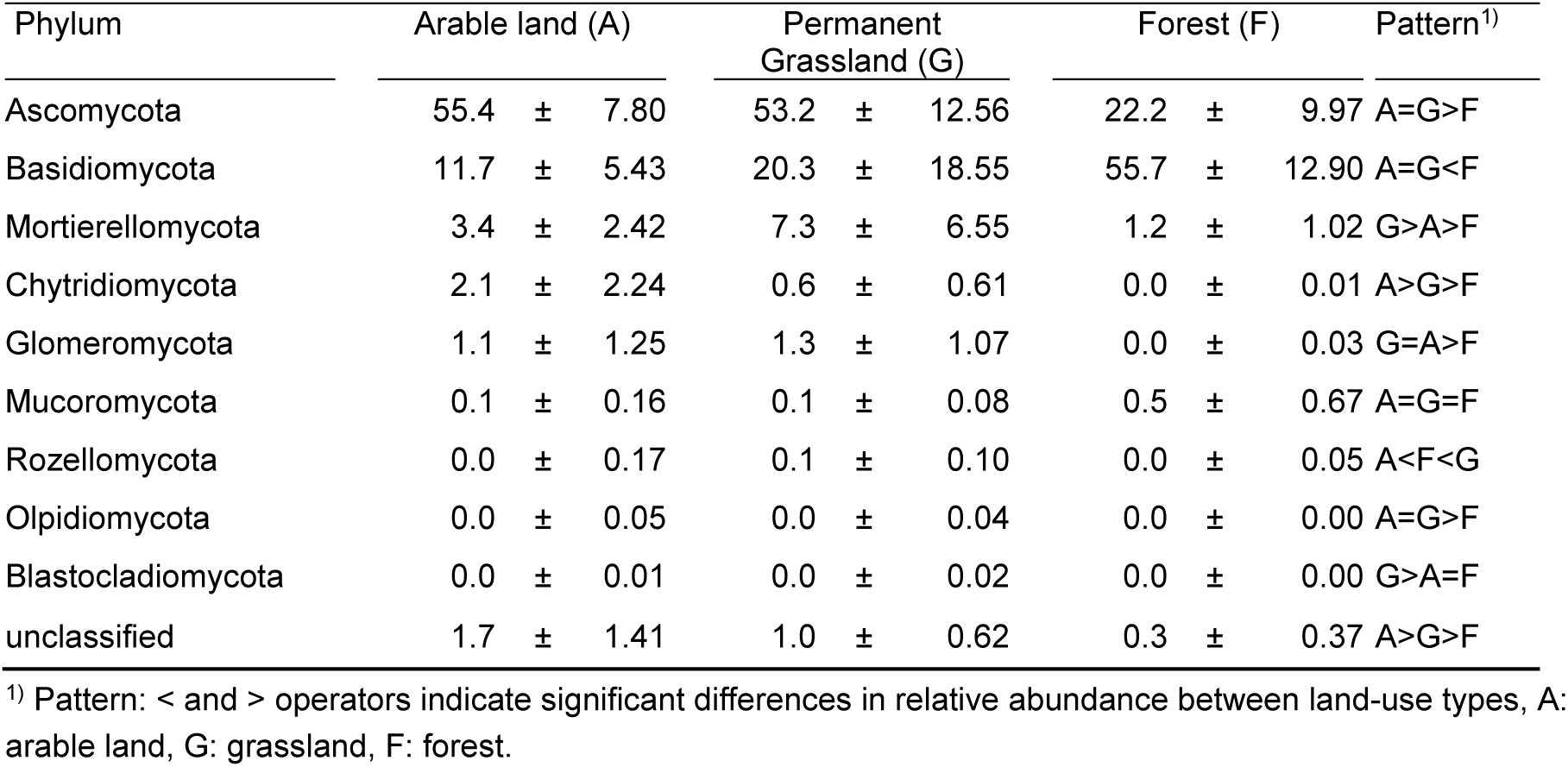
Relative abundances of fungal phyla in arable land, permanent grassland, and forest. Mean values (± standard deviations) of each land-use type are shown. Different letters indicate significant differences (Dunn Test, p < 0.05).

## References

Ai C, Zhang S, Zhang X et al. Distinct responses of soil bacterial and fungal communities to changes in fertilization regime and crop rotation. Geoderma 2018;319: 156–66.

Anderson M, Gorley RN, Clarke K. PERMANOVA+ for primer: Guide to software and statistical methods, 2008.

Anderson MJ. A new method for non-parametric multivariate analysis of variance. Austral Ecology 2001;26: 32–46.

Anderson MJ, Ellingsen KE, McArdle BH. Multivariate dispersion as a measure of beta diversity. Ecology Letters 2006;9: 683–93.

Anderson MJ, Willis TJ. Canonical analysis of principal coordinates: a useful method of constrained ordination for ecology. Ecology 2003;84: 511–25.

Babin D, Deubel A, Jacquiod S et al. Impact of long-term agricultural management practices on soil prokaryotic communities. Soil Biology and Biochemistry 2019;129: 17–28.

Bahnmann B, Mašínová T, Halvorsen R et al. Effects of oak, beech and spruce on the distribution and community structure of fungi in litter and soils across a temperate forest. Soil Biology and Biochemistry 2018;119: 162–73.

Bahram M, Hildebrand F, Forslund SK et al. Structure and function of the global topsoil microbiome. Nature 2018;560: 233–7.

Banerjee S, Walder F, Büchi L et al. Agricultural intensification reduces microbial network complexity and the abundance of keystone taxa in roots. ISME J 2019;13: 1722–36.

Bardgett RD, van der Putten WH. Belowground biodiversity and ecosystem functioning. Nature 2014;515: 505.

Beck S, Anderson IC, Drigo B et al. A soil fungal metacommunity perspective reveals stronger and more localised interactions above the tree line of an alpine/subalpine ecotone. Soil Biology and Biochemistry 2019;135: 1–9.

Benjamini Y, Hochberg Y. Controlling the false discovery rate: A practical and powerful approach to multiple testing. Journal of the Royal Statistical Society: Series B (Methodological) 1995;57: 289–300.

Bürgmann H, Pesaro M, Widmer F et al. A strategy for optimizing quality and quantity of DNA extracted from soil. Journal of Microbiological Methods 2001;45: 7–20.

Carini P. A “Cultural” renaissance: Genomics breathes new life into an old craft. mSystems 2019;4: e00092–19.

Cavaletti L, Monciardini P, Bamonte R et al. New lineage of filamentous, spore-forming, gram-positive bacteria from soil. Applied and Environmental Microbiology 2006;72: 4360–9.

Clarke KR, Warwick RM. Change in marine communities: an approach to statistical analysis and interpretation, 2nd edition. PRIMER-E: Plymouth, 2001.

Costa OYA, Raaijmakers JM, Kuramae EE. Microbial extracellular polymeric substances: Ecological function and impact on soil aggregation. Frontiers in Microbiology 2018;9.

Cui H, Sun W, Delgado-Baquerizo M et al. The effects of mowing and multi-level N fertilization on soil bacterial and fungal communities in a semiarid grassland are year-dependent. Soil Biology and Biochemistry 2020;151: 108040.

De Cáceres M, Legendre P. Associations between species and groups of sites: indices and statistical inference. Ecology 2009;90: 3566–74.

Degrune F, Theodorakopoulos N, Colinet G et al. Temporal dynamics of soil microbial communities below the seedbed under two contrasting tillage regimes. Frontiers in Microbiology 2017;8.

Delgado-Baquerizo M. Obscure soil microbes and where to find them. ISME J 2019;13: 2120–4.

Delgado-Baquerizo M, Oliverio AM, Brewer TE et al. A global atlas of the dominant bacteria found in soil. Science 2018;359: 320–5.

Dequiedt S, Saby NPA, Lelievre M et al. Biogeographical patterns of soil molecular microbial biomass as influenced by soil characteristics and management. Global Ecology and Biogeography 2011;20: 641–52.

Edgar RC. Search and clustering orders of magnitude faster than BLAST. Bioinformatics 2010;26: 2460–1.

Egidi E, Delgado-Baquerizo M, Plett JM et al. A few Ascomycota taxa dominate soil fungal communities worldwide. Nature Communications 2019;10: 2369.

Fox A, Lüscher A, Widmer F. Plant species identity drives soil microbial community structures that persist under a following crop. Ecology and Evolution 2020;10: 8652–68.

Frey B, Carnol M, Dharmarajah A et al. Only minor changes in the soil microbiome of a sub-alpine forest after 20 years of moderately increased nitrogen loads. Frontiers in Forests and Global Change 2020;3.

Frey B, Rime T, Phillips M et al. Microbial diversity in European alpine permafrost and active layers. FEMS Microbiology Ecology 2016;92.

Frey B, Walthert L, Perez-Mon C et al. Deep soil layers of drought-exposed forests harbor poorly known bacterial and fungal communities. Frontiers in Microbiology 2021;12: 1061.

Gilmullina A, Rumpel C, Blagodatskaya E et al. Management of grasslands by mowing versus grazing – impacts on soil organic matter quality and microbial functioning. Applied Soil Ecology 2020;156: 103701.

Giraldo A, Crous PW. Inside Plectosphaerellaceae. Studies in Mycology 2019;92: 227–86.

Glassman SI, Martiny JBH. Broadscale ecological patterns are robust to use of exact sequence variants versus operational taxonomic units. mSphere 2018;3: e00148–18.

Glassman SI, Wang IJ, Bruns TD. Environmental filtering by pH and soil nutrients drives community assembly in fungi at fine spatial scales. Molecular Ecology 2017;26: 6960–73.

Griffiths RI, Thomson BC, James P et al. The bacterial biogeography of British soils. Environmental Microbiology 2011;13: 1642–54.

Griffiths RI, Thomson BC, Plassart P et al. Mapping and validating predictions of soil bacterial biodiversity using European and national scale datasets. Applied Soil Ecology 2016;97: 61–8.

Gschwend F, Aregger K, Gramlich A et al. Periodic waterlogging consistently shapes agricultural soil microbiomes by promoting specific taxa. Applied Soil Ecology 2020;155: 103623.

Gschwend F, Braun-Kiewnick A, Widmer F et al. Apple blossoms from a Swiss orchard with low-input plant protection regime reveal high abundance of potential fire blight antagonists. Phytobiomes Journal 2021;0: null.

Gschwend F, Hartmann M, Hug A et al. Long-term stability of soil bacterial and fungal community structures revealed in their abundant and rare fractions.

Gubler A, Wächter D, Schwab P et al. Twenty-five years of observations of soil organic carbon in Swiss croplands showing stability overall but with some divergent trends. Environmental Monitoring and Assessment 2019;191: 277.

Guerra CA, Heintz-Buschart A, Sikorski J et al. Blind spots in global soil biodiversity and ecosystem function research. Nature Communications 2020;11: 3870.

Hallin S, Philippot L, Löffler FE et al. Genomics and ecology of novel N2O-reducing microorganisms. Trends in Microbiology 2018;26: 43–55.

Hamilton NE, Ferry M. ggtern: Ternary diagrams using ggplot2. Journal of Statistical Software 2018;87: 1–17.

Hartmann M, Brunner I, Hagedorn F et al. A decade of irrigation transforms the soil microbiome of a semi-arid pine forest. Molecular Ecology 2017;26: 1190–206.

Hartmann M, Frey B, Kölliker R et al. Semi-automated genetic analyses of soil microbial communities: comparison of T-RFLP and RISA based on descriptive and discriminative statistical approaches. Journal of Microbiological Methods 2005;61: 349–60.

Hartmann M, Frey B, Mayer J et al. Distinct soil microbial diversity under long-term organic and conventional farming. ISME J 2015;9: 1177–94.

Hartmann M, Niklaus PA, Zimmermann S et al. Resistance and resilience of the forest soil microbiome to logging-associated compaction. ISME J 2014;8: 226.

Hemkemeyer M, Christensen BT, Tebbe CC et al. Taxon-specific fungal preference for distinct soil particle size fractions. European Journal of Soil Biology 2019;94: 103103.

Hemkemeyer M, Dohrmann AB, Christensen BT et al. Bacterial preferences for specific soil particle size fractions revealed by community analyses. Frontiers in Microbiology 2018;9.

Jiao S, Xu Y, Zhang J et al. Core microbiota in agricultural soils and their potential associations with nutrient cycling. mSystems 2019;4: e00313–18.

Joergensen RG. The fumigation-extraction method to estimate soil microbial biomass: Calibration of the kEC value. Soil Biology and Biochemistry 1996;28: 25–31.

Jones RT, Robeson MS, Lauber CL et al. A comprehensive survey of soil acidobacterial diversity using pyrosequencing and clone library analyses. ISME J 2009;3: 442–53.

Kaiser K, Wemheuer B, Korolkow V et al. Driving forces of soil bacterial community structure, diversity, and function in temperate grasslands and forests. Scientific Reports 2016;6: 33696.

Karimi B, Terrat S, Dequiedt S et al. Biogeography of soil bacteria and archaea across France. Science Advances 2018;4: eaat1808.

Kielak AM, Barreto CC, Kowalchuk GA et al. The ecology of Acidobacteria: Moving beyond genes and genomes. Frontiers in Microbiology 2016;7.

Kindt R, Coe R. Tree diversity analysis: a manual and software for common statistical methods for ecological and biodiversity studies. Nairobi: World Agroforestry Centre, 2005.

Klosterman SJ, Atallah ZK, Vallad GE et al. Diversity, pathogenicity, and management of *Verticillium* species. Annual Review of Phytopathology 2009;47: 39–62.

Kuhn M. Building predictive models in R using the caret package. Journal of statistical software 2008;28: 1–26.

Lammel DR, Barth G, Ovaskainen O et al. Direct and indirect effects of a pH gradient bring insights into the mechanisms driving prokaryotic community structures. Microbiome 2018;6: 106.

Lauber CL, Hamady M, Knight R et al. Pyrosequencing-based sssessment of doil pH as a predictor of soil bacterial community dtructure at the continental scale. Applied and Environmental Microbiology 2009;75: 5111–20.

Lazzaro A, Hartmann M, Blaser P et al. Bacterial community structure and activity in different Cd-treated forest soils. FEMS Microbiology Ecology 2006;58: 278–92.

Lee MR, Hawkes CV. Plant and soil drivers of whole-plant microbiomes: variation in switchgrass fungi from coastal to mountain sites. Phytobiomes Journal 2020, DOI 10.1094/PBIOMES-07-20-0056-FI: PBIOMES-07-20-0056-FI.

Leeuwen JPv, Saby NPA, Jones A et al. Gap assessment in current soil monitoring networks across Europe for measuring soil functions. Environmental Research Letters 2017;12: 124007.

Leff JW, Bardgett RD, Wilkinson A et al. Predicting the structure of soil communities from plant community taxonomy, phylogeny, and traits. ISME J 2018;12: 1794–805.

Liu XZ, Wang QM, Göker M et al. Towards an integrated phylogenetic classification of the Tremellomycetes. Studies in Mycology 2015;81: 85–147.

Mayerhofer J, Eckard S, Hartmann M et al. Assessing effects of the entomopathogenic fungus Metarhizium brunneum on soil microbial communities in *Agriotes* spp. biological pest control. FEMS Microbiology Ecology 2017;93.

Mayerhofer J, Wächter D, Calanca P et al. Environmental and anthropogenic factors shape major bacterial community types across the complex mountain landscape of Switzerland. Frontiers in microbiology 2021;12: 500.

McArdle BH, Anderson MJ. Fitting multivariate models to community data: a comment on distance-based redundancy analysis. Ecology 2001;82: 290–7.

Nacke H, Thürmer A, Wollherr A et al. Pyrosequencing-based assessment of bacterial community structure along different management types in German forest and grassland soils. PLOS ONE 2011;6: e17000.

Nilsson RH, Larsson K-H, Taylor AF S et al. The UNITE database for molecular identification of fungi: handling dark taxa and parallel taxonomic classifications. Nucleic Acids Research 2018;47: D259–D64.

Peralta AL, Sun Y, McDaniel MD et al. Crop rotational diversity increases disease suppressive capacity of soil microbiomes. Ecosphere 2018;9: e02235.

Piazza G, Ercoli L, Nuti M et al. Interaction between conservation tillage and nitrogen gertilization shapes prokaryotic and fungal diversity at different soil depths: evidence from a 23-year field experiment in the mediterranean area. Frontiers in Microbiology 2019;10.

Quan Y, Muggia L, Moreno LF et al. A re-evaluation of the Chaetothyriales using criteria of comparative biology. Fungal Diversity 2020;103: 47–85.

Quast C, Pruesse E, Yilmaz P et al. The SILVA ribosomal RNA gene database project: improved data processing and web-based tools. Nucleic Acids Research 2012;41: D590–D6.

R Core Team. R: A language and environment for statistical computing [Computer software manual]. Vienna, Austria. URL https://www.R-project.org/. 2016.

Rice AV, Currah RS. *Oidiodendron*: A survey of the named species and related anamorphs of Myxotrichum. Studies in Mycology 2005;53: 83–120.

Rivera-Becerril F, van Tuinen D, Chatagnier O et al. Impact of a pesticide cocktail (fenhexamid, folpel, deltamethrin) on the abundance of Glomeromycota in two agricultural soils. Science of The Total Environment 2017;577: 84–93.

RStudio. RStudio: integrated development for R. RStudio, Inc, Boston, MA *URL* http://www.rstudio.com 2015;42: 14.

Schloss PD, Westcott SL, Ryabin T et al. Introducing mothur: open-source, platform-independent, community-supported software for describing and comparing microbial communities. Applied and Environmental Microbiology 2009;75: 7537–41.

Steen AD, Crits-Christoph A, Carini P et al. High proportions of bacteria and archaea across most biomes remain uncultured. ISME J 2019;13: 3126–30.

Talbot JM, Bruns TD, Taylor JW et al. Endemism and functional convergence across the North American soil mycobiome. Proceedings of the National Academy of Sciences 2014;111: 6341–6.

Tedersoo L, Bahram M, Põlme S et al. Global diversity and geography of soil fungi. Science 2014;346: 1256688.

Tedersoo L, Bahram M, Puusepp R et al. Novel soil-inhabiting clades fill gaps in the fungal tree of life. Microbiome 2017;5: 42.

Vance ED, Brookes PC, Jenkinson DS. An extraction method for measuring soil microbial biomass C. Soil Biology and Biochemistry 1987;19: 703–7.

Větrovský T, Kohout P, Kopecký M et al. A meta-analysis of global fungal distribution reveals climate-driven patterns. Nature Communications 2019;10: 5142.

Walsh CM, Gebert MJ, Delgado-Baquerizo M et al. A global survey of mycobacterial diversity in soil. Applied and Environmental Microbiology 2019;85: e01180–19.

Wisnoski NI, Lennon JT. Microbial community assembly in a multi-layer dendritic metacommunity. Oecologia 2021;195: 13–24.

